# DHX30 coordinates cytoplasmic translation and mitochondrial function contributing to cancer cell survival

**DOI:** 10.1101/2020.07.13.196709

**Authors:** Bartolomeo Bosco, Annalisa Rossi, Dario Rizzotto, Sebastiano Giorgetta, Alicia Perzolli, Francesco Bonollo, Angeline Gaucherot, Frédéric Catez, Jean-Jacques Diaz, Erik Dassi, Alberto Inga

## Abstract

DHX30 was recently implicated in the translation control of mRNAs involved in p53-dependent apoptosis. Here we show that DHX30 exhibits a more general function by integrating the activities of its cytoplasmic isoform and of the more abundant mitochondrial one. The depletion of both DHX30 isoforms in HCT116 cells leads to constitutive changes in polysome-associated mRNAs, enhancing the translation of mRNAs coding for cytoplasmic ribosomal proteins while reducing the translational efficiency of the nuclear-encoded mitoribosome mRNAs. Furthermore, depletion of both DHX30 isoforms exhibits higher global translation but slower proliferation, and reduced mitochondrial energy metabolism. Isoform-specific silencing established a role for cytoplasmic DHX30 in modulating global translation. The impact on global translation and proliferation were confirmed in U2OS and MCF7 cells, although the effect of DHX30 depletion on mitochondrial gene expression was observed only in MCF7 cells. Exploiting RIP, eCLIP, and gene expression data, we identified a gene signature comprising DHX30 and fourteen mitoribosome transcripts that we candidate as direct targets: this signature shows prognostic value in several TCGA cancer types, with higher expression associated with reduced overall survival. We propose that DHX30 contributes to cell homeostasis by coordinating ribosome biogenesis, global translation, and mitochondrial metabolism. Targeting DHX30 could, thus, expose a vulnerability in cancer cells.

**Author summary:** Translation occurs in the cell both through cytoplasmic and mitochondrial ribosomes, respectively translating mRNAs encoded by the nuclear and the mitochondrial genome. Here we found that DHX30, an RNA-binding protein implicated in p53-dependent apoptosis, enhances the translation of mRNAs coding for cytoplasmic ribosomal proteins while reducing that of the mitoribosome mRNAs when silenced. This coordination of the cytoplasmic and mitochondrial translation machineries affected both cell proliferation and energy metabolism, suggesting an important role for this mechanism in determining the fitness of cancer cells. Indeed, the analysis of publicly available cancer datasets led us to define a 15-genes signature that is able to affect the prognosis of a subset of cancer types. In this subset, we found that higher expression of the genes composing the signature is associated with a worse prognosis. We thus propose DHX30 as a potential vulnerability in cancer cells, that could be targeted to develop novel therapeutic strategies.

## Introduction

The DHX30 RNA binding protein (RBP) is an ATP-dependent RNA helicase highly expressed in neural cells and somites during embryogenesis in mice. It plays an important role in development and its homozygous deletion is lethal for embryos [1]. In humans, DHX30 is involved in the antiviral function of the zinc-finger ZAP protein. Indeed, it was demonstrated that these two proteins directly interact with each other via the N-terminal domain and that DHX30 is necessary for the optimal antiviral activity of ZAP [2]. Recently, six de novo missense mutations were identified in DHX30 in twelve unrelated patients affected by global developmental delay (GDD), intellectual disability (ID), severe speech impairment and gait abnormalities [3]. All mutations caused amino acid changes in the highly conserved helicase motif, impairing the protein’s ATPase activity or RNA recognition. Moreover, overexpression of those DHX30 mutants led to increased propensity to trigger stress granules (SG) formation and decreased global translation [3]. It was shown that DHX30 could play an important role also in human fibroblast and osteosarcoma mitochondria. Indeed, in that compartment it interacts with a Fas-activated serine-threonine kinase (FASTKD2), modulating mitochondrial ribosome maturation and assembly [4].

Our group and collaborators recently compared p53-mediated responses to Nutlin-3 treatment in three different cancer cell lines: HCT116, SJSA1 and MCF7 [5,6]. It was observed that p53 activation caused cell cycle arrest in HCT116, massive apoptosis in SJSA1 and an intermediate phenotype in MCF7, consistent with earlier reports [7]. By analyzing the cell lines’ translatomes (i.e. transcripts loaded on polysomes and in active translation), we discovered that only 0.2% of the genes were commonly differentially expressed (DEGs). Furthermore, the polysome profiling of SJSA1 revealed an enrichment of pro-apoptotic transcripts [6] in the fraction corresponding to polysomes. Those featured instances of a specific cis-element in their 3’UTR, labeled as CGPD, standing for CG-rich motif for p53-dependent death. The DExH box RNA helicase DHX30 was found to be one RBP whose binding to the CGPD motif correlated with a specific cell fate. Indeed, DHX30 was expressed at higher levels in HCT116 than SJSA1, resulting in a lower translation of GCPD-motif-containing transcripts in HCT116 cells. Depletion of DHX30 in HCT116 by stable shRNA increased the translation potential of mRNAs containing the CGPD motif, resulting in higher levels of apoptosis after p53 activation by Nutlin treatment, mimicking in part the apoptotic phenotype of SJSA1 [6].

In this work, we set to determine the broader phenotypic role of DHX30 in the colorectal cancer cell line HCT116. By virtue of two alternative promoters leading to the production isoforms containing or not a mitochondrial localization signal, DHX30 can couple mitochondrial function, ribosome biogenesis and global translation. Depleting DHX30 in HCT116 cells reduced cell proliferation and sensitised cells to small molecules targeting mitochondrial function. Results were extended to two different cancer cell models and their implications explored using TGCA data.

## Results

### DHX30 depletion enhances translation in HCT116 cells

We previously identified DHX30 as a negative modulator of p53-dependent apoptosis, performing polysomal profiling followed by RNA-seq of HCT116 cells clones stably depleted for DHX30 and treated with the MDM2 inhibitor Nutlin-3 [7]. Here we reanalyzed this dataset (GSE95024), complementing it by performing a total-cytoplasmic RNA profiling of the same cells by RNA-seq (GSE154065), and focusing on the comparison between DHX30-depleted (shDHX30) and control (shNT) HCT116 cells in the untreated control.

We compared the gene expression changes at the polysomal or total cytoplasmic RNA levels and performed a gene set enrichment analysis [8]. This revealed an increased abundance of ribosomal proteins and ribosome biogenesis genes in shDHX30 cells, only at the polysomal level (**Figure 1A**, **Table S1**). Consistently, analysis of translation efficiency (TE), measured as the ratio between relative RNA abundance in polysomal and total cytoplasmic fractions, revealed that DHX30 depletion is associated with higher TE for transcripts coding for ribosomal protein subunits (RPL, RPS, **Figure 1B**). On the other end, nuclear transcripts coding mitochondrial ribosomal proteins exhibited reduced TE in DHX30-depleted cells, despite showing higher constitutive TE compared to the cytoplasmic ribosomal transcripts (mRPL, mRPS, **Figure 1B**). Validation by qPCR was performed for four RPs and four MRPs transcripts (**Figure S1A**). The basal differences in TE between ribosome and mitoribosome transcripts was confirmed in all cases, and for five of them results reproduced the effect of DHX30 depletion on TE that was observed from the RNA-seq data. A trend for higher expression of RPL and RPS transcripts and lower expression of MRPL and MRPS transcripts in DHX30-depleted cells was also apparent (**Figure S1B**).

**Figure 1.**
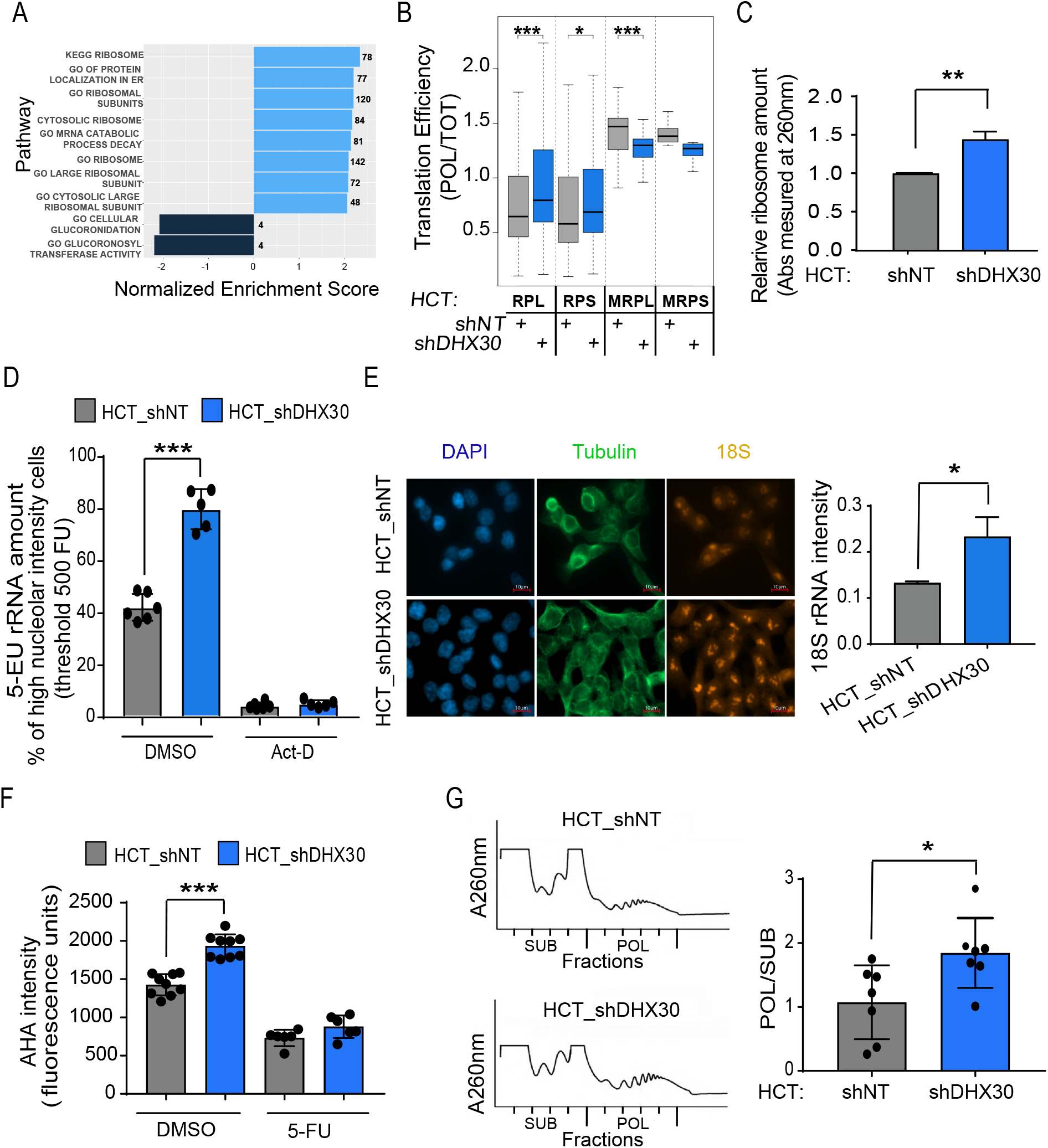
DHX30 reduces ribosome biogenesis and translation in HCT116 colorectal cancer cells. **A)** Most significant terms by Gene Set Enrichment Analysis of the polysome-bound DEGs (HCT_shDHX30 vs HCT sh_NT). Bars plot the Normalized Enrichment Score, (NES). Numbers indicate the number of genes in the leading edge. See Table S1 for full results. **B**) Box plot of Translation efficiency for the indicated transcript groups. The RNA-seq CPM in polysomal over total RNA was measured. Results obtained in HCT116_shDHX30 cells are compared to the shNT control clone; *p < 0.05, ***p < 0.001. **C**) Amounts of ribosomes produced in HCT116_shDHX30 and HCT_shNT. After ribosomes isolation, protein absorbance was measured at λ 260nm. Data are normalized on shNT and are mean ± SD (n=3); *p < 0.05. **D**) Amount of ribosomal RNA estimated based on the fluorescence intensity of 5-ethynyl uridine (5-EU) incorporated in nascent rRNAs present in the nucleoli. HCT116_shDHX30 cells are compared to the shNT control clone in untreated condition while Actinomycin-D treatment was used as a control. Data are mean ± SD (n=6); ***p < 0.001. **E**) (Left) Representative images of one of three independent experiments of Fluorescent In Situ Hybridization (FISH) for rRNA precursor 18S (red). Immunofluorescence of tubulin (green) and staining with DAPI (blue) were used to visualize cells. (Right) Box plot of 18S rRNA intensity quantification comparing HCT116_shDHX30 with HCT_shNT analyzed with Cell Profiler software. Data are expressed as mean per well ± SD of three biological replicates in which 10 nuclei were quantified for each replicate; *p < 0.05. **F**) Analysis of global translation based on the fluorescence intensity of L-azidohomoalanine (AHA) incorporated in nascent proteins present in the cytoplasm. HCT116_shDHX30 cells are compared to the shNT control clone in untreated condition while 5-Fluorouracil treatment was used as positive control. Data are mean ± SD (n=6 to 9); ***p < 0.005. **G**) (Left) Polysome profiling of HCT_shNT and HCT_shDHX30, revealed by the measurement of absorbance at a wavelength of 260 nm. (Right) Box plot of the relative quantification of the area under the curve of fractions corresponding to the polysomes related to the fractions corresponding to the ribosomal subunits and 80S monosome (POL/SUB, an estimate of translation efficiency). Data are mean ± SD (n=7); *p < 0.05.

We thus sought to understand if DHX30 could be directly regulating this class of genes and exploited the data of DHX30 eCLIP assay performed in K562 cells as part of the ENCODE project [9]. Despite the different cell line, the eCLIP data indicate that DHX30 can bind 67 ribosomal (36 RPLs and 31 RPSs) and 23 mitoribosomal (16 MRPLs and 7 MRPSs) protein transcripts (**Table S1**) [9].

Consistent with the higher TE for RPL and RPS transcripts, DHX30 depletion in HCT116 cells was associated with higher relative amounts of ribosomes extracted and quantified from a sucrose cushion (**Figure 1C**), and higher rRNA levels (**Figure 1D, E**). Furthermore, DHX30-depleted cells not only showed evidence of increased ribosome biogenesis, but also of higher global translation (**Figure 1F, 1G**). The higher global translation does not appear to be related to an increase in MYC expression or activity (**Figure S1C**), nor to a significant increase in the activity mTOR pathway (**Figure S1D**). Finally, DHX30 protein was found to be associated with ribosomal subunits, 80S monosomes as well as with low molecular-weight polysomes in HCT116, MCF7 and U2OS cells (**Figure S1F**).

Hence, DHX30 appears to negatively influence ribosome biogenesis, global translation and also the translation of specific mRNAs. It is important to emphasize that these are not necessarily dissociated events, as a change in ribosome number could be sufficient to impact on translation specificity [10].

### DHX30 depletion alters the expression of mitoribosomal transcripts

The impact of DHX30 depletion on the expression and translation efficiency of nuclear-encoded mitoribosomal proteins caught our attention, as it showed the opposite trend compared to ribosomal protein transcripts (**Figure 1B, Figure S1**). First, we established the potential for DHX30 to bind to mitoribosomal protein transcripts by RIP assays, choosing MRPL11 (UL11m), that was previously evaluated in cancer cells [3], and MRPS22 (mS22), as a representative of the large and small subunits, respectively (**Figure 2A**). Next, we observed that DHX30-depleted cells showed reduced expression of these two genes at both the RNA and protein levels (**Figure 2B, C**). These findings suggest that DHX30 directly promotes stability and/or translation of mitoribosome transcripts. The modulation of mitoribosome proteins expression might impact on mitochondrial translation and contribute to the functions described for DHX30 in the mitochondrial matrix [4,11]. Notably, the DHX30 gene contains two promoters, and the transcript resulting from the internal, more 3’-one (ENST00000457607.1) includes an alternative first exon that contains a predicted mitochondrial targeting sequence [12,13]. Mitochondrial localization of the protein was confirmed by immunofluorescence (**Figure 2D**), consistent with previous results in human fibroblasts [4] and APEX-seq data [14]. We checked the relative levels of the DHX30 transcripts from the two promoters and found the putative mitochondrial isoform to be about four times more abundant (**Figure 2E**), as indicated also by western blot analysis on extracts of mitochondrial and cytoplasmic fractions (**Figure 2F**).

**Figure 2.**
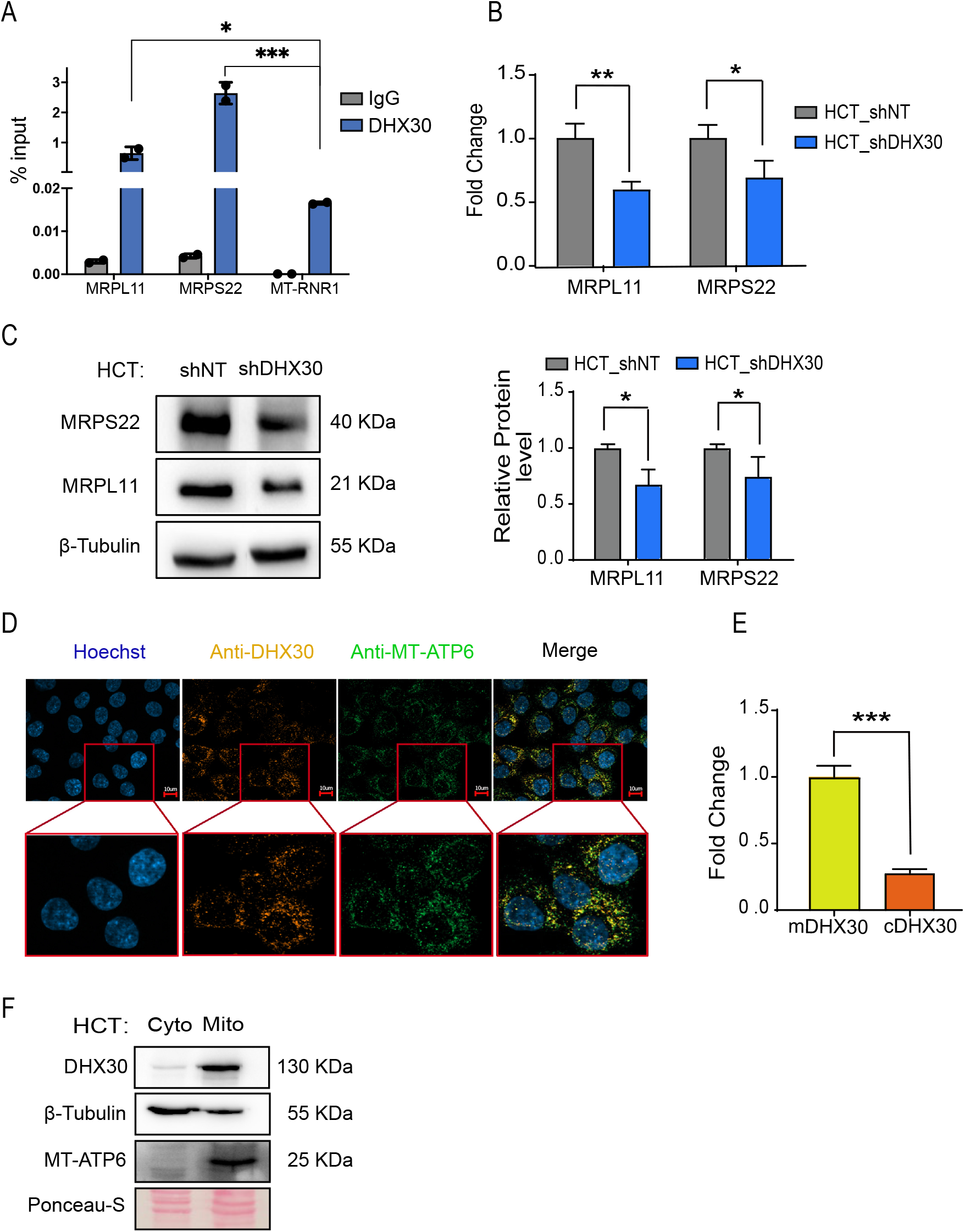
Cytoplasmic DHX30 modulates the expression of nuclear encoded mito-ribosome components MRPL11 and MPRS22. **A**) RNA immunoprecipitation experiments to study the binding of DHX30 to MRPL11 and MRPS22 transcripts. Results obtained with a primary antibody targeting DHX30 (blue) or the control IgG control (grey) are plotted relative as % of input. Data are mean and individual points (n=2); p-value was calculated comparing the amount of each sample with the amount of *RNR1* mRNA; *p < 0.05, ***p < 0.001. **B**) Relative mRNA levels of MRPL11 and MRPS22 in HCT116_shDHX30 compared to the shNT control clone. Data are mean ± SD (n=3); *p < 0.05; **p < 0.01. **C**) (Left) Protein levels of MRPL11 and MRPS22. β-Tubulin was used as loading control. Immunoblots represent one of three independent experiments. (Right) MRPL11 and MRPS22 protein quantification in HCT116_shDHX30 normalized on shNT control clone. Data are mean ± SD (n=3); *p < 0.05. **D**) Representative images of one of three independent immunofluorescence experiments to colocalize DHX30 (red) with MT-ATP6 (green, used as mitochondrial marker). Staining with Hoechst (blue) was used to visualize cells’ nuclei. **E**) Relative mRNA levels of cytoplasmic DHX30 (cDHX30) and the mitochondrial variant (mDHX30, set to 1). Data are mean ± SD (n=3); ***p < 0.001. **F**) Relative DHX30 protein levels in cytoplasm and mitochondria. β-Tubulin and MT-ATP6 are used as controls of cytoplasmic-mitochondrial fractionation and Ponceau-S staining is used as loading control. One of three independent experiments is shown.

Stable DHX30-silencing was obtained using three different shRNAs [6] that targeted both cytoplasmic and mitochondrial transcripts (from here on labeled cDHX30 and mDHX30) (**Figure S2A**), and isolating single clones. To confirm results, we also employed transient silencing using one siRNA specific for cDHX30 (labeled siDHX30-C) or an siRNA targeting transcripts from both promoters (labeled siDHX30-C+M). The activity and specificity of the two siRNAs was assessed by qRT-PCR 48 and 96 hours post-silencing (**Figure S2B**, **Figure 3A**). Transient silencing of just cDHX30 or both cDHX30 and mDHX30 for 96 hours led to an increase in global translation (**Figure 3B**) that was not associated with an increase in the activity mTOR pathway (**Figure S1E**), thus confirming the observations obtained with the different stable clones, but indicating that cDHX30 modulates global translation.

**Figure 3.**
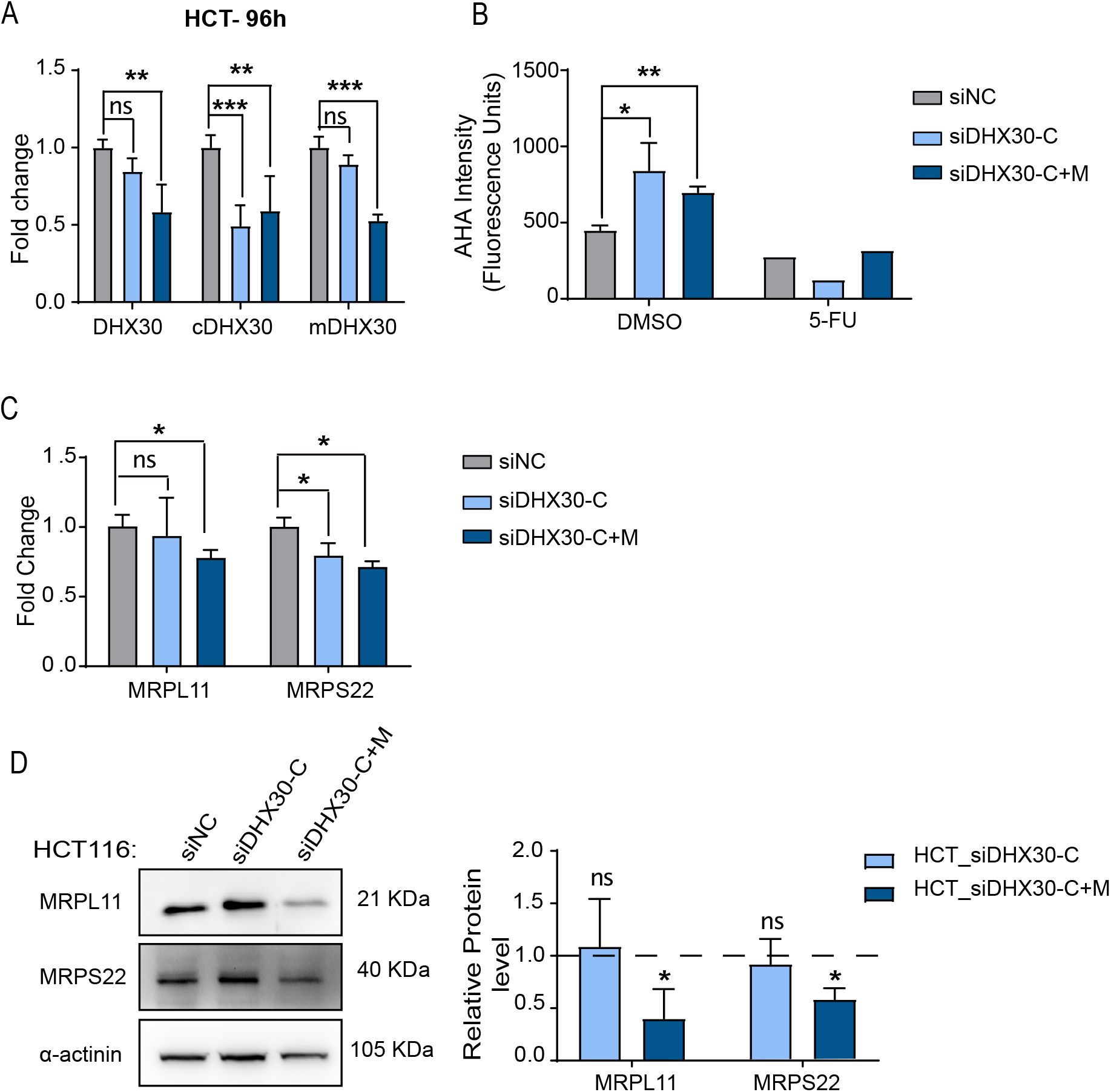
Transient DHX30 silencing leads to enhanced translation but reduced mitorobosome gene expression in HCT116 cells. **A)** qRT-PCR to verify the silencing of DHX30 transcripts in HCT116 96 hours after transient transfection with the indicated siRNAs. The relative mRNA levels of cytoplasmic (cDHX30) and mitochondrial (mDHX30) DHX30 transcripts as well as of total DHX30 (measured with a pair or primers common to all annotated transcripts). Data are mean ± SD (n=3); **p < 0.01; ***p < 0.001. **B)** Global translation based on the fluorescence intensity of L-azidohomoalanine (AHA) incorporated in nascent proteins present in the cytoplasm. HCT116 were transiently transfected with the indicated siRNAs and tested after 96 hours. 5-Fluorouracil treatment was used as control treatment leading to translation inhibition. Data are mean ± SD (n=3); * p < 0.05; **p < 0.01. **C**) mRNA levels of MRPL11 and MRPS22 in HCT116 silenced transiently for cytoplasmic DHX30 (HCT_siDHX30-C) or for both cytoplasmic and mitochondrial variants (HCT_siDHX30-C+M). Data are compared to the siRNA negative control (HCT_siNC) and are mean ± SD (n=3); *p < 0.05. **D**) (Left) MRPL11 and MRPS22 protein levels in HCT116 silenced for cytoplasmic DHX30 (HCT_siDHX30-C) or for both cytoplasmic and mitochondrial variants (HCT_siDHX30-C+M). Immunoblot represents one of three independent experiments. (Right) MRPL11 and MRPS22 protein quantification in HCT116_siDHX30-C and HCT116_siDHX30-C+M normalized on α-actinin and compared to siNC control. Data are mean ± SD (n=3); *p < 0.05.

Transient silencing of both cDHX30 and mDHX30 resulted in a reduction of the expression of MRPL11 and MRPS22 (**Figure 3C, D**) similar to what we previously observed with stable depletion of both isoforms (Figure 2), while cDHX30 silencing was ineffective, considering the results at both RNA and protein levels.

DHX30 stable or transient silencing was extended to U2OS, osteosarcoma-derived cells that express relatively high levels of DHX30 (**Figure S2C, S2D**). Stable depletion of both cDHX30 and mDHX30 was associated with a significant reduction of MRPL11 (RNA and protein) and of MRPS22 mRNA expression (**Figure S2E, S2F**), while transient depletion by siRNAs did not significantly impact on the expression of the two mitoribosome transcripts (**Figure S2G**), although MRPL11 protein levels were reduced by siDHX30-C+M (**Figure S2H**).

Transient silencing of both cDHX30 and mDHX30 was also performed in the breast cancer-derived MCF7 cells (**Figure S3A**) and led to a reduction in the expression of both MRPL11 and MRPS22 mRNAs (**Figure S3B**), although only for MRPL11 the effect was consistent also at protein level (**Figure S3C**). Cytoplasmic DHX30 transient silencing in U2OS and MCF7 also led to higher global translation, confirming the results obtained in HCT116 cells (**Figure S3D**, **S3E**).

### DHX30 depletion impacts mitochondrial gene expression and function

We next focused on the expression of mitochondrially encoded genes. First of all, we established that DHX30 depletion did not impact on the number of mitochondrial genomes by digital PCR in HCT116 cells (**Figure 4A**). Instead, the expression of several mitochondrially encoded transcripts was significanly reduced (**Figure 4B**). The steady-state expression of two mitochondrially encoded proteins, MT-ATP6 and MT-ATP8, was investigated and shown to be reduced in HCT116_shDHX30 cells (**Figure 4C, S3F**). We pursued the same analysis after transient silencing, confirming the lower expression of mitochondrially encoded genes at the RNA and protein levels, but only when the DHX30 mitochondrial variant (siDHX30-C+M) was silenced and analysed 96 hours post silencing (**Figure 4D, 4E**). In fact, although DHX30 depletion was visible already after 48 hours (**Figure S2B**) from the addition of the siRNAs, more time was needed to appreciate a reduction in the expression of mitochondrially encoded genes. The expression of mitochondrially encoded genes was checked also after DHX30 silencing in U2OS and MCF7 cells (**Figure S4**). In U2OS_shDHX30 cells, all of the tested transcripts, with the exception of MT-ND6, were down-modulated compared to the U2OS_shNT control (**Figure S4A,** left panel). Instead, none of the seven tested transcripts were down-modulated by transient DHX30 silencing, regardless of the siRNA used (**Figure S4A,** right panel). In fact, three of them were slightly upregulated. This latter result is consistent with the observation of a slightly lower efficacy of DHX30 depletion by siRNAs and with the lack of an impact on MRPL11 or MRPS22 expression (Figure S2). MT-ATP6 was also examined by western blot and its amount was reduced in U2OS_shDHX30 cells (**Figure S4C**), but not in U2OS cells that were transiently silenced for DHX30 (**Figure S4D**). Transient DHX30 depletion (by siDHX30 C+M) in MCF7 cells led to a significant reduction in the expression of all tested mitochondrial transcripts, although for MT-ATP6 the reduction was not significant at protein levels (**Figure S4B, S4E**).

**Figure 4.**
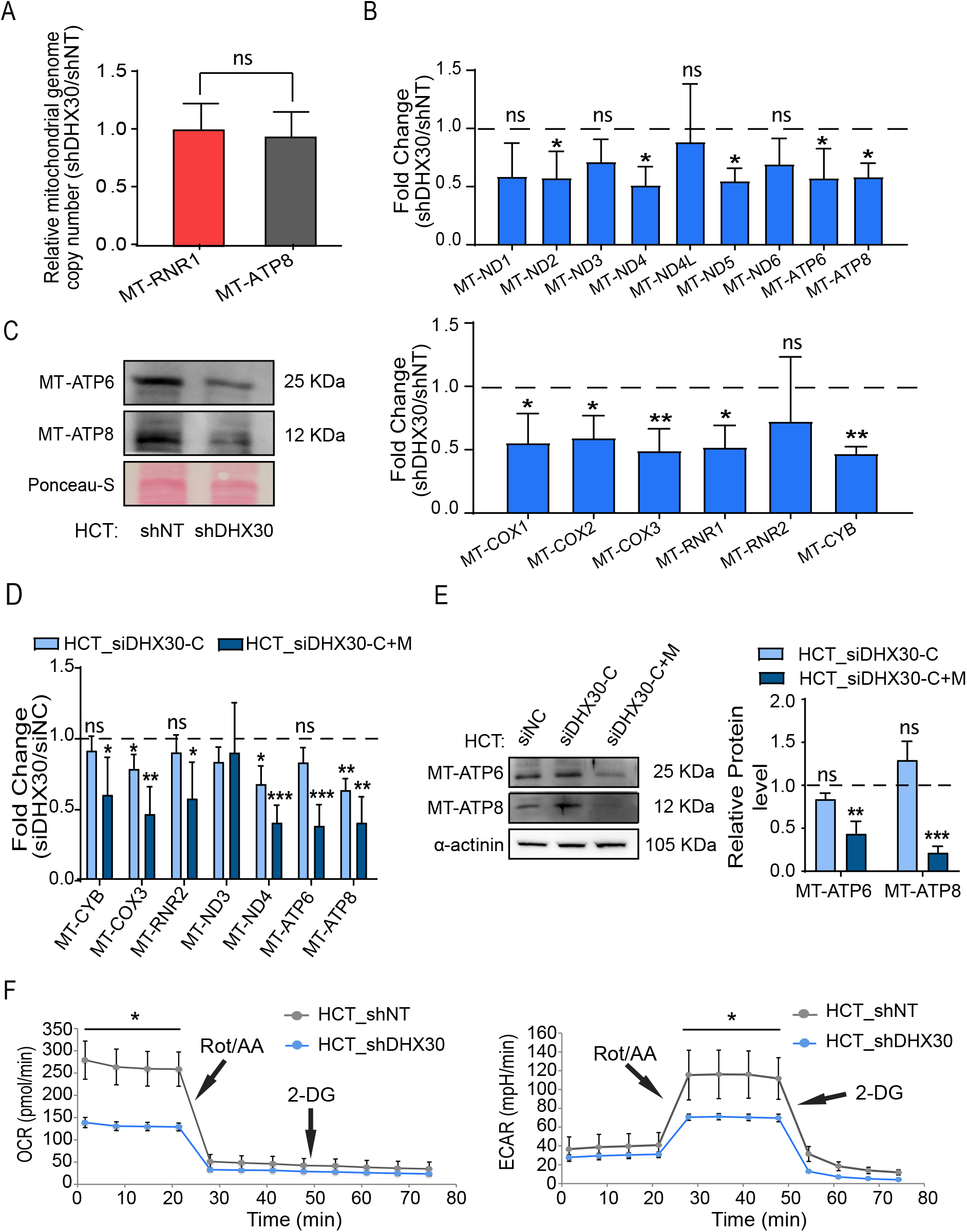
Depletion of DHX30 reduces the expression and function of mitochondrially encoded OXPHOS components. **A**) Relative mitochondrial genome copy number in HCT116_shDHX30 and -shNT measured by droplet digital PCR. MT-RNR1 and MT-ATP8 were amplified along with the nuclear diploid marker gene CTDSP1. Bars plot mean ± SD (n=3). **B**) Relative mRNA levels of mitochondrially encoded OXPHOS components in HCT116_shDHX30 compared to HCT116_shNT (dashed line, set to 1). For both upper and lower panel, data are mean ± SD (n=3); *p < 0.05; **p < 0.01. **C**) Protein levels of MT-ATP6 and MT-ATP8 in mitochondrial lysates of HCT116_shDHX30 and the shNT control clone. Ponceau-S was used as loading control. Immunoblots represent one of two independent qualitative comparisons. **D**) Relative mRNA levels of the indicated mitochondrially encoded OXPHOS components in HCT116 transiently silenced for DHX30 expression (siDHX30-C, siDHX30-C+M) compared to siNC (dashed line, set to 1). Data are mean ± SD (n=3); *p < 0.05; **p < 0.01; ***p < 0.001. **E**) (Left) Protein levels of MT-ATP6 and MT-ATP8 in HCT116 transiently silenced as in D). α-actinin was used as loading control. Immunoblots represent one of three independent experiments. (Right) MT-ATP6 and MT-ATP8 protein quantification in HCT116_siDHX30-C and HCT116_siDHX30-C+M normalized on α-actinin and compared to siNC control. Data are mean ± SD (n=3); **p < 0.01; ***p < 0.001. **F**) (Left) Measurement of the oxygen consumption rate (OCR) to evaluate mitochondrial respiration by Seahorse XF analyzer. Points before the Rotenone/Antimycin-A (Rot/AA) treatment correspond to the basal mitochondrial respiration. (Right) Extracellular acidification rate (ECAR) measurement was used as a means to measure glycolysis. Points before and after the Rotenone/Antimycin-A (Rot/AA) treatment correspond respectively to basal and compensatory glycolysis -in response to the block of mitochondrial respiration-. 2-Deoxyglucose (2-DG) is then used to block glycolysis. For both panels, one of three independent replicates is presented. Data are mean ± SD (n=3 wells in the Seahorse cartridge); *p < 0.05.

Finally, we evaluated markers of carbon metabolism and mitochondrial respiration capacity by an Agilent SeahorseXF real-time analyzer. HCT116_shDHX30 cells had a significantly lower basal oxygen consumption rate compared to the control cell line. Notably, DHX30-depleted cells did not show compensatory glycolysis in basal culture condition, not even when we chemically abolished mitochondrial respiration. We indeed observed a lower compensatory glycolysis compared with the control (**Figure 4F**). These results suggest that HCT116_shDHX30 cells do not show a typical Warburg effect [15], and are expected to produce less energy due to reduced mitochondrial respiration.

Collectively, the depletion of DHX30 seems to generate an imbalance in cell homeostasis, with higher demand of chemical energy from ribosome biogenesis and cytoplasmic translation but lower mitochondrial activity.

### DHX30 depletion impaired cell proliferation rate and increased apoptosis proneness

Next, we characterized the phenotypic impact of DHX30 depletion in HCT116. Consistent with the lower mitochondrial oxygen consumption rate, HCT116_shDHX30 cells showed a significantly lower proliferation rate compared to the control cell clone. This was observed by colony formation assay and confirmed through time-course proliferation by cell count via high-content microscopy or using a Real-Time Cell Analyzer xCELLigence (**Figure 5A-C**). Interestingly, we observed a reduction in proliferation after DHX30 silencing also in a spheroid assay (**Figure 5D**). The lower growth rate in DHX30 depleted cells cannot be attributed to an arrest in a particular phase of the cell cycle (**Figure 5E**). Instead, it could be associated with a moderate increase in the phosphorylation of eIF2alpha (**Figure 5F**), but not to an obvious ER-stress response, based on the expression of CHOP, DDIT4, TRIB3, and on the XBP1 splicing pattern (**Figure S5A**).

**Figure 5.**
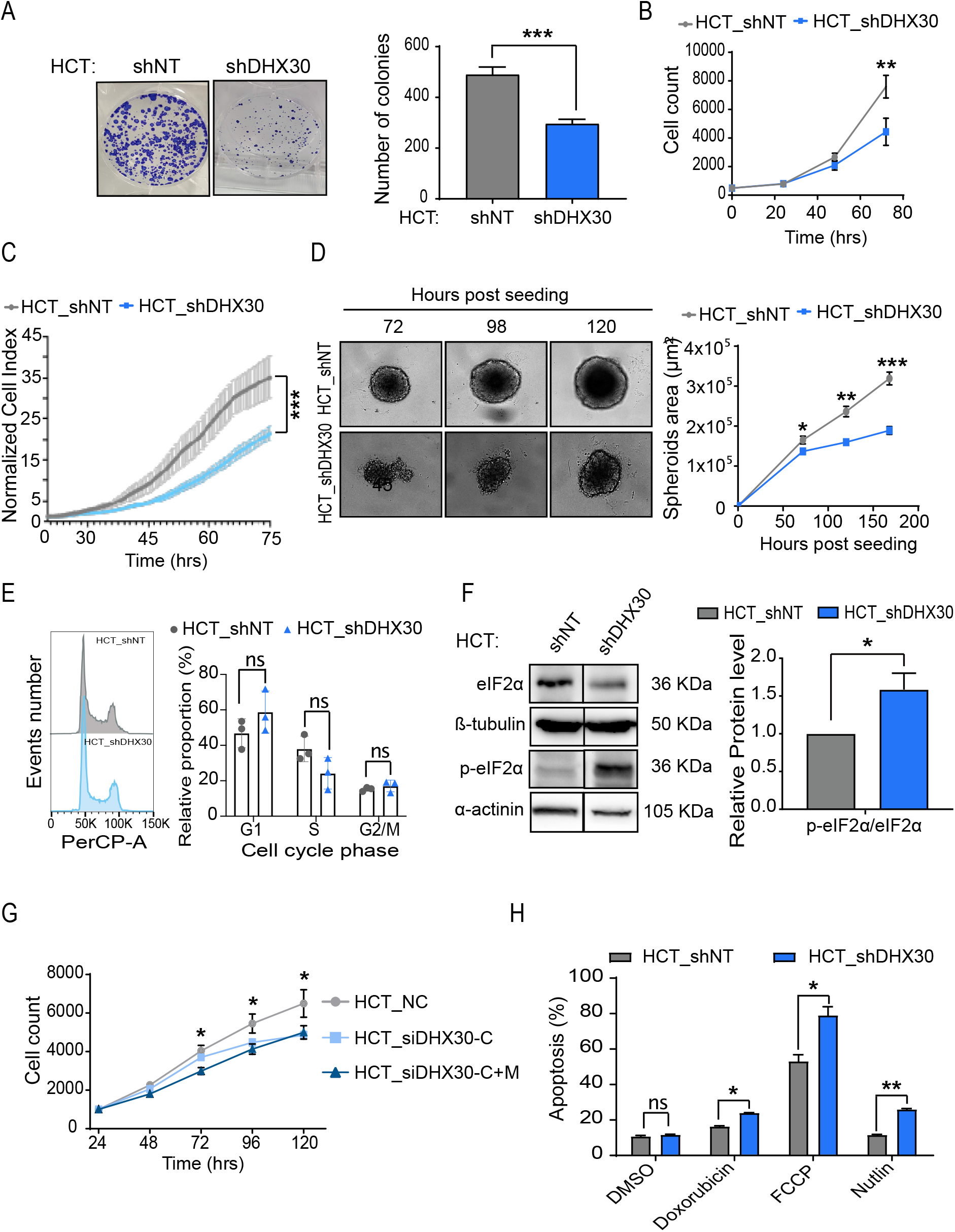
Depletion of DHX30 reduces proliferation and survival in basal and treatment conditions. **A)** (Left) Representative image of colony formation assays. (Right) Colony quantification by ImageJ software. Data are mean ± SD (n=3). **B)** Relative cell proliferation measured by high-content microscopy in digital phase contrast. Data are mean ± SD (n=3). **C)** Estimate of relative proliferation by an impedance-based Real-Time Cell Analyzer. Data are mean ± SD (n=3). **D)** Spheroid formation and growth assay. (Left) A representative image at the indicated time points is shown in the left panel. (Right) Spheroid area measured by ImageJ software. Data are mean ± SD (n=3). **E)** (Left) Representative image of HCT116_shNT and shDHX30 cell cycle profiles. (Right) Quantification of cell cycle profile in HCT116 DHX30 depleted cells compared to shNT control. Data are mean ± SD (n=3). **F**) Lef panel: relative expression of eIF2alpha and phosho-eIF2alpha in HCT116 cells stably depleted for DHX30 expression or control. A representative immunodetection image is shown. Right panel: relative average protein quantification and SD of three replicates is presented in the bar graph; n = 3, *p < 0.05. **G)** Relative proliferation measured as in B) but starting 24 hours after transient silencing DHX30 with the indicated siRNAs. **H**) Relative expression of the annexin-V apoptosis markers in cells treated for 48 hours with the indicated drugs or DMSO control. For all panels, data are mean ± SD (n=3); *p < 0.05; **p < 0.01.

A reduction in proliferation was confirmed also by cell counting upon transient silencing of cDHX30 and, particularly, of both cDHX30 and mDHX30 (**Figure 5G**). The impact on cell proliferation was also examined using U2OS cells stably depleted for DHX30 that showed a significant reduction of proliferation compared to the U2OS_shNT control (**Figure S5B**). Transient silencing did not impact on the proliferation of U2OS cells, consistent with the results observed on DHX30 target gene expression, while it led to a significant reduction in MCF7 cells (**Figure S5B**).

Furthermore, DHX30 depletion was associated with increased apoptosis after 48 hours of Nutlin treatment, as expected by our previous results [6], but also when cells were treated with the topoisomerase inhibitor doxorubicin and with FCCP, an agent that causes mitochondrial membrane depolarization (**Figure 5H**).

### DHX30 signature and cancer outcome

Our results suggest that DHX30 exerts a constitutive function that improves cellular fitness by balancing energy metabolism and global translation potential. Furthermore, our previous study identified DHX30 as a negative modulator of the translation of specific mRNAs, thus controlling p53-dependent apoptosis [6]. Both these functions suggest that DHX30 could be a modifier of cancer cell properties potentially impacting on clinical variables.

Although total RNA-seq data are not a good proxy for investigating translation controls [16], in this study we showed that DHX30 depletion impacts on steady-state levels of nuclear-encoded mitoribosomal transcripts. MRPL11 and, particularly, MRPS22 can be considered direct DHX30 targets. We next cross-referenced our TE and GSEA data with the RIP results and DHX30 eCLIP data in ENCODE [9] and compiled a list of 14 mitoribosomal protein (MRP) transcripts considered DHX30 direct target candidates (**Figure 6**, **Figure S6**). Interrogating RNA-seq data of TGCA tumors through the GEPIA web resource [17,18], the expression of DHX30, and of each MRP transcript or of the group of 14 candidate target MRPs appeared to be positively correlated in several cancer types (**Figure 6, Figure S6**), including adrenocortical carcinoma, and hepatocellular carcinomas. No such correlation was instead apparent for other cancer types, including breast or colon adenocarcinomas. We next used this gene signature to stratify patients’ clinical outcome. Interestingly, for the cancer types where a positive correlation in the expression of DHX30 and the 14 MRPs was observed, the combined 15-gene signature (*i.e.* including DHX30) showed a prognostic value. In particular, higher expression was associated with reduced Overall Survival or Disease-Free Survival (**Figure 6**). Instead, the comparison of DHX30 expression between tumors and matched controls did not show a consistent trend but confirmed a rather wide variation among samples, and tissue-specific effects including both cases of over-expression and down-regulation (**Figure S7**). Finally, a pan-tissue view revealed a general positive correlation between the expression of ribosome and mitoribosome protein transcripts in normal samples, that is lost in cancer (**Figure S8**). These preliminary analyses suggest that a functional signature could be developed to predict aggressiveness in cancer types where DHX30 appears to stimulate mitoribosomal proteins expression.

**Figure 6.**
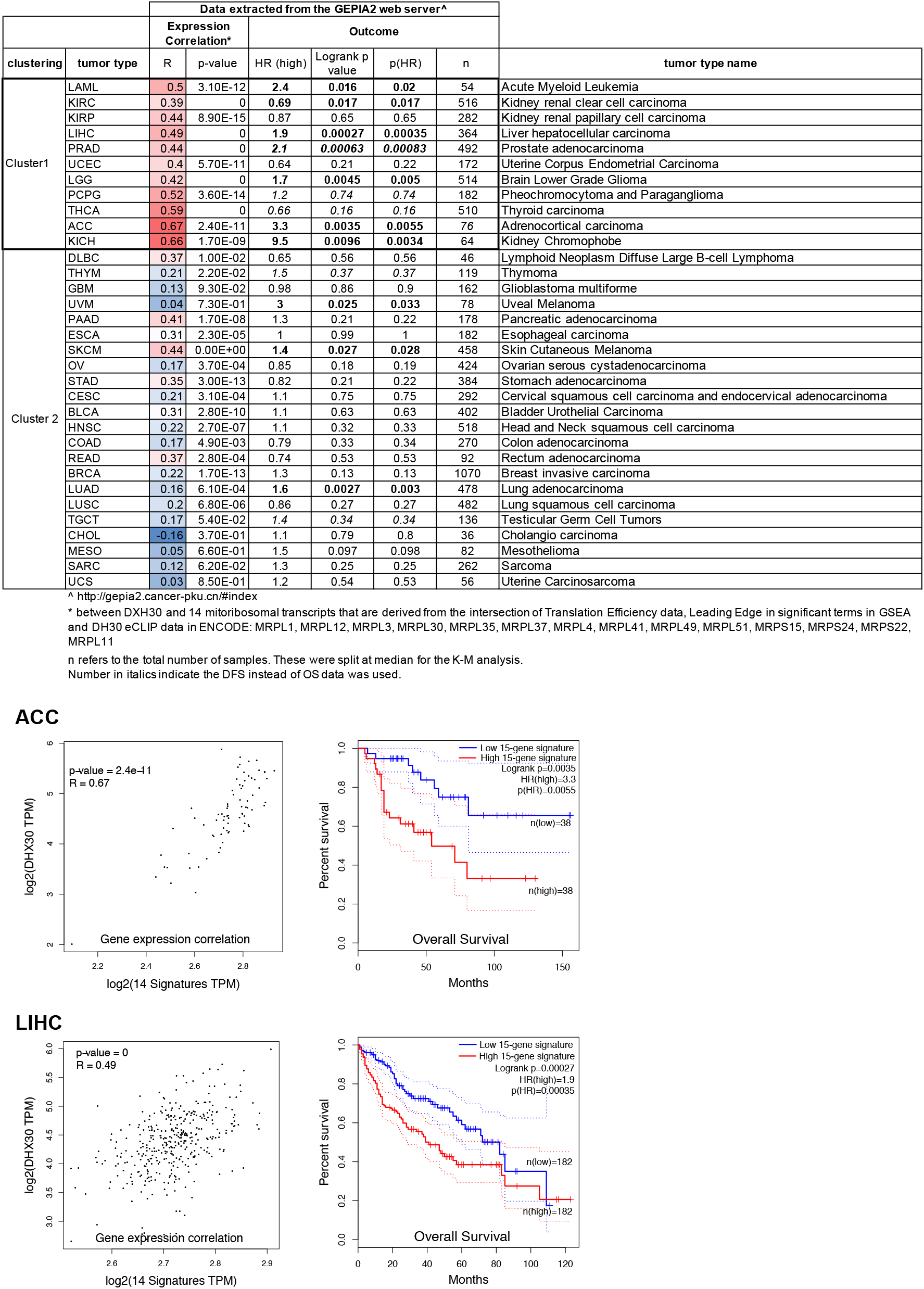
DHX30 expression positively correlates with the expression of mitoribosomal protein transcripts and can have prognostic significance. The table presents the expression correlation data (R value and p value) between DHX30 and the combined group of 14 mitoribosomal protein transcripts is listed for each cancer type (see also Figure S6). Data from Kaplan-Meier survival analysis for the aggregated 15-gene signature (DHX30 + the 14 mitoribosomal protein transcripts) are shown. Significant differences are highlighted in bold font. The tumor type acronym and extended name are also shown. The graphs below the table presents two examples among the cluster of eleven cancer types where a positive correlation is apparent. Left panels: Pearson correlation. Right panels: Overall Survival Kaplan-Meier. The correlation value, the Hazardous Ratio, p values and sample sizes are shown.

## Discussion

Ribosome biogenesis and translation impose a high metabolic demand on the cell [19,20]. Hence, a coordination between translational control and metabolic output ultimately involving mitochondrial respiratory functions is expected to contribute to cell homeostasis and fitness [21,22]. However, relatively few proteins and pathways have been established to exert a direct role in balancing cytoplasmic translation initiation with mitochondrial metabolism [23,24]. eIF6 represents a significant example, as it has been clearly shown that it negatively controls 80S monosome assembly, a necessary step for translation initiation, while at the same time playing a critical positive role in mitochondrial functions, as revealed by broad changes in the mitochondrial proteome in eIF6 hemizygous mice [25–27].

We propose that DHX30 can also exert an important housekeeping role in coordinating ribosome biogenesis, translation, and mitochondrial respiration. DHX30 depletion in HCT116, stable or transient, leads to a modest but significant increase in ribosome biogenesis as well as in global translation; on the contrary, mito-ribosome transcripts, particularly of the large subunits, exhibit reduced translation efficiency. Furthermore, DHX30 can also exert a more direct role on mitochondria, as the protein can directly localize to the organelle, thanks to an alternative first exon that features a localization signal. The steady state expression of several mitochondrially encoded genes was reduced following the depletion of both DHX30 transcripts and perhaps more effectively, by the reduction of the mitochondrial isoform. A previous study provided clear evidence that DHX30 together with DDX28, FASTKD2, and FASTKD5 can promote the assembly of 55S mito-monosome and translation [4]. Consistent with the relevance of a mitochondrial and translation function, the DHX30 transcript has been shown to be located and locally translated at the ER-Outer Mitochondrial membrane interface, by APEX-seq [14]. In fact, none of the DHX30 closer homologs showed strong evidence of such localized translation, or other evidence of mitochondrial localization.

A previous report also showed evidence for DHX30 interaction with mitochondrial transcripts in human fibroblasts by RIP-seq [4]. Our data instead point to a direct interaction with mitoribosome transcripts and their positive modulation as another means by which DHX30 can indirectly affect mitochondrial translation. We validated by RIP the binding of DHX30 to MRPL11 and MRPS22. Leveraging public data from eCLIP experiments in ENCODE and our polysomal profiling, RNA-seq, and GSEA analysis, we propose that DHX30 could directly interact with fourteen mito-ribosome protein transcripts. An even larger set of ribosomal protein transcripts are nominated as direct DHX30 targets by eCLIP [9] and are showing changes in translation efficiency upon DHX30 depletion in HCT116 cells.

We observed that these two groupings of transcripts markedly differed for their basal translation efficiency, which was low for cytoplasmic ribosomal protein transcripts. Those mRNAs are known to contain particularly structured 5’-UTR and to be strongly regulated at the level of translation initiation by mTOR and MYC-regulated pathways [28–30]. The mechanism by which DHX30 can control ribosomal protein (RP) transcripts remains to be established. In a recent study, we discovered an RNA sequence motif in 3’UTRs, labeled CGPD, that is targeted by DHX30 and can mediate higher translation levels [6]. While we cannot exclude that the CGPD motif could be implicated, only a subset of RP transcripts harbors instances of it. Our *de novo* search for over-represented motifs did not retrieve another prominent candidate *cis*-element. An even lower number of mito-RP transcripts were found to harbor instances of the CGPD-motif. This was expected, as DHX30 is inferred to have an opposite role on RP and mito-RP transcripts based on the direction of changes in their TE observed in HCT116 shDHX30 cells. For the mito-RP gene group, a *de novo* motif over-representation search did not identify a strongly enriched cis-element.

The magnitude of fold-change and translation efficiency changes for both ribosomal and mitoribosomal transcripts in response to DHX30 silencing is modest in absolute value. This could be in part due to a limitation of the experimental approach. As reported in our earlier study, a complete knock-out of DHX30 does not seem to be attainable in HCT116 cells. Both our siRNA and even more so the shRNA clones retain partial DHX30 expression. Furthermore, it is important to emphasize that changes in the expression or even polysomal association of ribosomal protein transcripts do not necessarily predict translation rate changes. However, several endpoints consistently suggested that HCT116 shDHX30 cells exhibited higher ribosome number and increased global translation. When treated with an MDM2 inhibitor inducing strong p53-dependent transcriptional response, those cells were also shown to markedly alter their translatome [6]. Our results pointed to a direct role of DHX30 on translation specificity. However, we cannot exclude that the global effect on ribosome biogenesis we propose represents a constitutive, housekeeping function for DHX30. This function would also contribute to the observed changes in translation specificity, under the notion that a change in global translation potential or in the modulation of translation initiation is not expected to impact equally every available transcript in a cell [10,22].

Depletion of DHX30 reduced cell proliferation in various assays. This effect was visible also by transient silencing of just the cytoplasmic transcript, although it was more evident when both cytoplasmic and mitochondrial isoforms were targeted. Although we cannot exclude that the lower proliferation results from a checkpoint activation, as we have seen that in the siRNA experiments the lack of p53 can reduce the proliferation lag (**Figure S9**), cells did not show evidence for overt cell cycle arrest. This is not entirely unexpected, due to the residual levels of DHX30 expression in the cell models we used, as noted above, and also for the possible selective pressure for compensatory effects given the central nature of the processes involved. Transient silencing of DHX30 in HCT116 p53 null cells besides having a reduced impact on proliferation also did not significantly alter the expression of MRPS22, MRPL11 or of mitochondrial transcripts. While this may be suggestive of a functional interaction between DHX30 and p53, results may be affected by the lower efficiency of DHX30 depletion in HCT116 p53^−/−^ cells (**Figure S9**).

As several other RBPs, mito-ribosomal proteins have been proposed to have additional, moonlighting functions unrelated to mito-ribosome biogenesis, in particular in the modulation of apoptosis. For example, MRPS29, also known as DAP3, was reported to influence the extrinsic apoptosis pathway [31,32], while MRPL41 was proposed to modulate p53-dependent intrinsic apoptosis [33]. It is, however, unlikely that these potential pro-apoptotic functions can contribute to the proliferation defects seen after DHX30 depletion, given that their expression is not significantly modified. The expression of several mito-ribosome components has been evaluated as potential biomarkers associated with cancer clinical variables [34].

Hence, we reasoned that changes in DHX30 levels to an extent that would not lead to overt stress responses could provide opportunities for increased fitness due to the coordination between mitochondrial function and translation potential, which could be a balancing function between energy supply and demand. If any, the effects would be expected to reflect tissue-specific differences in metabolism. We explored TGCA data, first evaluating if a positive correlation could be observed in normal tissue and/or primary cancer samples for DHX30 and mito-ribosome gene expression. Cross-referencing eCLIP data, with our polysomal profiling, RNA-seq, RIP results, and GSEA data we compiled a list of fourteen mitoribosomal transcripts as candidate direct DHX30 targets. Interestingly DHX30 and this 14-gene list showed a prognostic significance in cancer types where a positive correlation in their expression was observed. Instead, copy number alterations or mutations in DHX30 were a rare event in cancer (8.6%), were not associated with specific tissue or cancer types, and, although based on a few cases, seemed to be mutually exclusive with mitochondrial genome changes (**Figure S10**).

In conclusion, our study demonstrates the role of DHX30 in the coordination between global cytoplasmic translation and mitochondrial functions. This role contributes to the oncogenic potential of cancer cells, and appears to correlate with tumor aggressiveness and clinical outcome. Furthermore, during stress responses activating p53, DHX30 can reduce the apoptotic commitment of cancer cells by acting on specific pro-apoptotic transcripts, thus providing a potential actionable target for therapeutic purposes [6].

## Materials and Methods

### RNA-seq library preparation and sequencing

shDHX30 and control cells were either kept untreated or treated with 10μM Nutlin, and processed after 12h. Polysomal profiling and RNA extraction of reconstituted total cytoplasmic fractions was performed as recently described [6]. Sequencing libraries were obtained following the manufacturer instructions of the TruSeq RNA Library preparation kit v2. We used 1.5 μg of RNA as input, and assessed the input RNA quality by the Agilent RNA 6000 Nano kit on a Agilent 2100 Bioanalyzer instrument. Four replicates were generated for each condition. The resulting 16 samples were sequenced on a HiSeq 2500 machine producing ~25M raw reads per each sample.

### RNA-seq data analysis

Reads were first quality filtered and trimmed with trimmomatic (minimum quality 30, minimum length 36nt) [35]. Then, each Gencode v27 (http://www.gencodegenes.org/releases/) transcript was quantified with Salmon [36]. Eventually, edgeR [37] was used to call Differentially Expressed Genes (DEGs) between conditions at the polysomal and total levels (shDHX30 DMSO vs shNT DMSO), using a 0.05 threshold on the adjusted p-value. GSEA was performed with the fgsea R package [8], including the Hallmark, Canonical Pathways, and GO gene sets. We used 1000 permutations to compute the significance p-value and used a BH adjusted p-value threshold of 0.05. Translational efficiency (TE) was computed on normalized expression data (counts per million reads, CPM) as the ratio of polysomal CPMs over total CPMs for each replicate and transcript in the annotation. Differences in TE (between conditions and groups of genes) and between polysomal and total samples were assessed by the Wilcoxon test. Motifs in ribosomal and mitoribosomal genes 5’- and 3’-UTRs were obtained with DREME [38] using a 1.0E-04 p-value threshold and shuffled input sequences as controls.

### Cell lines and culture conditions

HCT116 and U2OS shNT control or shDHX30 clones were obtained as recently described [6]. Briefly, cells were transduced with lentiviral vectors containing a pLK0.1 plasmid expressing shRNAs. DHX30 targeting sequences are: TRCN0000052028 (GCACACAAATGGACCGAAGAA) TRCN0000052031 (CCGATGGCTGACGTATTTCAT), TRCN0000052032 (GAGTTGTTTG ACGCAGCCAAA). Stable clones were selected exploiting the puromycin resistance marker. Single clone isolates were obtained and characterized for DHX30 protein depletion. Consistent results were obtained using the three different shRNAs HCT116 p53^−/−^ cells were obtained from the Vogelstein lab (Sur S. et al, 2009). MCF7 cells were purchased from ICLC (IRCCS San Martino Hospital, Genoa, Italy). All cell lines were cultured in RPMI (Gibco, Thermo Fisher Scientific, Waltham, MA, USA) supplemented with 10% Fetal Bovine Serum (Gibco, Thermo Fisher Scientific), 1X L-Glutamine and Pen/Strep (Gibco, Thermo Fisher Scientific) at 37 °C with 5% CO2 in a humidified atmosphere. U2OS cells were cultured in DMEM pH 7.4 (Gibco, Thermo Fisher Scientific) supplemented with 10% Fetal Bovine Serum (Gibco, Thermo Fisher Scientific), 1X L-Glutamine and Pen/Strep (Gibco, Thermo Fisher Scientific) at 37 °C with 5% CO2 in a humidified atmosphere. For routine culture of depleted clones and controls, 0.2 μg/mL Puromycin was added to the culture medium to maintain selection of the vectors. Puromycin was removed 24 hours before starting a specific experiment to avoid confounding effects. MCF-7, U2OS and HCT116 parental and p53^−/−^ cells were silenced for cytoplasmic or cytoplasmic + mitochondrial DHX30 variants using 25nM siRNAs (Trifecta, IDT, Coralville, IA, USA) (**Table S2**) transfected with Interferin (Polyplus-transfection, Illkirch, France). All experiments were performed at least 24 hours post-silencing.

### Western Blot

Cells were cultured in 6-well tissue culture plates. Cells were collected with trypsin-EDTA after 24 hours, centrifuged and washed with PBS. Samples were then lysed with RIPA buffer and the proteins were quantified by BCA assay (EUROCLONE, Milan, Italy). 30 μg of extracted proteins were loaded on 12% or 15% Tris-glycine Gel and then transferred onto a nitrocellulose membrane using Tris-Glycine buffer. Blocking was performed overnight with 5% not-fat dry milk, 0.1% TWEEN and PBS 1X. Immunodetection was obtained using primary and secondary antibodies reported in **Table S3**. Membranes were analyzed by ECL and detected with ChemiDocTM XRS+ (Bio-Rad, Hercules, CA, USA) using ImageLab software (Bio-Rad).

### Polysome profiling

Polyribosome analysis was performed as described in [39–41]. Briefly, HCT116 cells were grown on 15 cm Petri dishes with standard media and serum. When the cells reached 80% of confluence, cycloheximide (0.01 mg/ml, Sigma Aldrich) was added and kept in incubation for 10 min. Then cells were washed two times with cold PBS containing cycloheximide (0.01 mg/ml) and lysed with the following lysis buffer: 20 mM Tris-HCl (pH 7.5), 100 mM KCl, 5 mM MgCl2, 0.5% Nonidet P-40, 100 U/ml RNase inhibitors. Mitochondria and nuclei-free lysates were loaded onto 15–50% (w/v) density sucrose gradients in salt solution (100 mM NaCl, 5 mM MgCl2, 20 mM Tris–HCl pH 7.5), and ultracentrifuged at 180 000 × g for 100 min at 4°C. The sedimentation profiles were monitored by absorbance at 254 nm using a Teledyne ISCO UA-6 fractionator coupled to UV detector, collecting thirteen 1 ml fractions. RNA from pooled polysomal fraction was extracted and processed as described in [6]. For Western blot analysis, 100 ul of 100% TCA and 1 ml of ice-cold acetone were added to 1 ml of each fraction. Then samples were put at −80°C overnight to induce protein precipitation. Subsequently samples were centrifuged at 16,000 × g for 10 min at 4 °C and washed three times with 1 mL of ice-cold acetone. Finally, the pellets were solubilized directly in Laemmli buffer pH 8.

### Ribosome isolation

Ribosome isolation was performed following a protocol previously described [42]. Briefly, 8 × 10^6^ cells were seeded in a T150 flask, 48 hours before the procedure. At 80% of confluence, cells were detached and counted. 1 × 10^7^ cells were pelleted by 500 x g centrifugation for 5 minutes at 4°C and washed once with cold PBS. Supernatant was discarded and the pellet was resuspended gently with 300μL of cold buffer A (sucrose 250mM, KCl 250mM, MgCl2 5mM, and Tris-Cl 50 mM pH 7.4), added in three sequential steps, pipetting after every addition. To perform cell lysis, an appropriate volume of NP-40 was added to the homogenized cellular solution, in order to obtain a 0.7% (v/v) final concentration of the detergent, and cells were incubated on ice for 15 minutes, homogenizing the suspension by gentle pipetting every 5 minutes. Then the cell lysate was centrifuged at 750 x g for 10 minutes at 4°C to pellet nuclei; the recovered cytoplasmic fraction (supernatant) was centrifuged again at 12.500 x g for 10 minutes at 4°C to obtain a mitochondria pellet. Supernatant containing ribosomes was collected and its volume was accurately measured using a graduated pipet. KCl 4M solution was then slowly added in order to reach a final concentration of 0.5M. Meanwhile, 1mL of the sucrose cushion solution (sucrose 1M, KCl 0.5M, MgCl2 5mM and Tris-Cl 50mM pH 7.4) was added into 3-mL polycarbonate tube for an ultracentrifuge TL100.3 rotor (Beckman Coulter, Brea, CA, USA). The KCl-adjusted ribosome-containing solution was carefully added above the sucrose cushion and tubes were balanced by weight using buffer B (sucrose 250mM, KCl 0.5M, MgCl2 5mM and Tris-Cl 50mM pH 7.4). Then, they were ultracentrifuged at 250.000 x g for 2 hours at 4°C. At the end of the centrifugation, the supernatant was discarded and a very compact and dense translucent pellet containing ribosomes was quickly rinsed twice by carefully adding 200μL of cold water. Then, pellet was resuspended in 300μL of buffer C (KCl 25mM, MgCl2 5mM and Tris-Cl 50mM pH 7.4), adding the solution in three sequential steps and gently homogenized by pipetting after every 100μL addition. Finally, to estimate the amounts of ribosomes, absorbance at 260nm of the suspensions was measured by a nanodrop spectrophotometer (ThermoFisher).

### Global Translation

Global translation assay was performed using Click-iT^®^ AHA Alexa Fluor^®^ 488 Protein Synthesis HCS Assay (Thermo Fisher Scientific). Briefly, HCT116 shNT and shDHX30 cells were cultured in 96-well tissue culture plate and treated with 100mM 5-Fluorouracil (Sigma Aldrich) as positive control. After 24 hours of treatment, the culture medium was removed and 50μM L-azidohomoalanine prepared in pre-warmed L-methionine-free medium (Lonza) was added and cells were incubated for 1 hour. Cell fixation was performed using 3.7% formaldehyde (Sigma Aldrich) incubating the plate at room temperature for 15 minutes. After two washing with 3% BSA in PBS 1X cells were permeabilized with 0.5% Triton^®^ X-100 (Sigma Aldrich) and incubated for 15 minutes at room temperature. Cells were washed twice with 3% BSA in PBS 1X and Click-iT^®^ reaction cocktail was added incubating for 30 minutes at room temperature, protected from light. After the incubation reaction cocktail was removed, cells were washed twice with 3% BSA in PBS 1X and stained with 5μg/mL Hoechst (Sigma Aldrich). Samples’ images were acquired and analysed by high content fluorescence microscope Operetta^®^ (PerkinElmer, Waltham, MA, USA) using the following filter set: Alexa Fluor^®^ 488: Ex495/Em519 nm; Hoechst: Ex350/Em461 nm.

### Fluorescent In Situ Hybridization (FISH) on rRNA precursors

Cells were cultured on a coverslip place onto a 48-well plate and after 48 hours they were washed with PBS and fixed with 4% Paraformaldehyde (PFA) (Sigma Aldrich) for 30 minutes at RT. Then, cells were washed twice with PBS 1X and incubated in 70% ethanol overnight at 4°C. The day after, cells were rehydrated with 10% formamide in 2X SSC (Sigma Aldrich), twice for 5 minutes at RT. Meanwhile, buffer A [5μL formamide, 2.5μL SSC 2X, 2.5μL tRNA (10 ng/μL), water up to 20.25μL and then 2.5μL of each probe (10 ng/μL)] was incubated for 5 minutes at 90°C and then mixed quickly together with buffer B [25μL Dextran sulfate 20% in SSC 4X, 1.25μL Bovine Serum Albumin (BSA) (10ng/μL) and 2.5μL ribonucleoside vanadyl complex (RVC) (200mM)]. Then, 50μL A+B probe solution was dropped onto the coverslip and it was incubated in a humidified chamber for 3 hours at 37°C. When the incubation was finished, the coverslip was washed twice with 10% formamide in SSC 2X for 30 minutes at RT and once with PBS 1X for 5 minutes. Cells were stained with a solution of 0.5μg/mL DAPI (Sigma Aldrich) in PBS 1X for 5 minutes, washed three times with PBS 1X and the coverslip was finally placed with mounting medium on glass slide. Finally, images were acquired using Zeiss Observer z1 fluorescent microscope (Carl Zeiss, Oberokochen, Germany) with ZEN 2 blue edition software ver. 2.3 (Carl Zeiss). Images’ fluorescence intensity was measured using Cell Profiler software ver. 3.1.9 (Broad Institute, Cambridge, MA, USA).

### rRNA biogenesis

rRNA biogenesis was performed following the protocol described by [43,44]. Briefly, cells were cultured in 96-well tissue culture plates and after 24 hours were treated with Actinomycin D 100nM (Sigma Aldrich) as positive control. After 24 hours of treatment, 1mM 5-Ethynyluridine (Sigma Aldrich) was added to the culture medium and the plate was incubated for 2 hours. This short incubation with 5-EU allows a higher rate of incorporation in rRNAs than mRNAs, reducing the background signal. Then, cells were fixed using a working solution 1 (WR1) composed by 125mM Pipes pH 6.8 (Sigma Aldrich), 10mM EGTA (Sigma Aldrich), 1mM MgCl2 (Merck Millipore, Burlington, MA, USA), 0.2% Triton® X-100 (Sigma Aldrich) and 3.7% formaldehyde (Sigma Aldrich). Cells were washed twice with TBS 1X and stained for 30 minutes at room temperature using a working solution 2 (WR2) composed by 100mM Tris-HCl pH 8.5 (Sigma Aldrich), 1mM CuSO4 (Merck Millipore), 10μM fluorescent azide (Sigma Aldrich) and 100mM ascorbic acid (Merck Millipore). After four washes with TBS containing 0.5% Triton X-100, cells were stained with 0.5μg/mL Hoechst (Sigma Aldrich) and samples’ images were acquired and analysed by high content fluorescent microscope Operetta® (PerkinElmer) using the following filter set: Alexa Fluor® 488: Ex495/Em519 nm; Hoechst: Ex350/Em461 nm and setting a threshold on Alexa Fluor® 488 at 500 fluorescent units to plot the percentage of cells with high nucleolar signal intensity.

### Colony formation

1.0×10^3^ HCT116 shNT and shDHX30 cells per well were seeded in a 6-well plate and incubated. Culture medium was changed every three days. Colonies were ex fixed with 3.7% formaldehyde and stained with a solution containing 30% Methylene blue (Sigma Aldrich) in water, then washed five times for 5 minutes. Images were acquired and analysed using ImageJ software 1.8.0 (National Institute of Health, Bethesda, MD, USA).

### Cell Count in High-Content analysis and Real Time Cell Index Analysis

To analyze cell proliferation by cell count, 1.0×10^3^ cells per well were seeded in 96-well plates and incubated. Images were acquired by high content fluorescent microscope Operetta^®^ (PerkinElmer) in digital phase contrast for three days. To analyse cell proliferation by Real Time Cell Analyser xCELLigence^®^ (RTCA) (Roche, Basel, Switzerland) the background was set adding the only cell culture medium in each well of E-plate and incubating 30 minutes. After incubation, background was read. Then 1.0×10^3^ cells per well were seeded and incubated for 30 minutes and then the E-plate was inserted in the RTCA and impedance-based cell proliferation was estimated by RTCA readings every 15 minutes in the course of three days.

### Spheroid assay formation

3.0×10^3^ HCT116 shNT and shDHX30 cells per well were seeded in U-bottom ultra-low attachment 96-well plate (Corning Incorporated, Corning, NY, USA), centrifuged at 3000 rpm for 5 minutes and incubated. Starting after three days of incubation, images were acquired every 24 hours for eight additional days by microscope (Leica, Wetzlar, Germany). Spheroids’ area was measured by ImageJ software.

### Immunofluorescence

3.0×10^5^ cells per well were seeded on glass coverslip inserted in 6-well plates and incubated overnight. The days after, cells were fixed in 4% Paraformaldehyde (PFA) for 15 min. Then the PFA solution was removed and cells were rinsed twice with PBS 1X. After washing, cells were incubated for 1 hour with blocking solution (PBS 1X, 5% BSA and 0.5% Triton X-100). Next, cells were incubated overnight at 4°C with primary antibodies (Table 2.4) diluted in blocking solution. The following day, after three washes with PBS 1X, the anti-mouse and anti-rabbit secondary antibodies conjugated with AlexaFluor 488 or AlexaFluor 594 (Invitrogen, Carlsbad, CA, USA) (diluted 1:1000) were added to the samples and incubated for 1 hour at RT under agitation. Cells were washed with PBS 1X three times and cell nuclei were stained with 0.5μg/mL Hoechst (Sigma Aldrich) (diluted 1:5000). Finally, coverslips were mounted on glass slides and images were acquired using Zeiss Observer Z1 fluorescent microscope and Zen 2012*®* software (Carl Zeiss Jena, Germany). The list of primary antibodies used are reported in **Table S3**.

### Compensatory Glycolytic Test

3.0×10^3^ cells per well were seeded in Xfp cell culture microplate (Agilent Technologies, Santa Clara, CA, USA) and incubated overnight while XF calibrant solution was added in each well of extracellular flux cartridge and incubated overnight without CO2. The day after, culture media was removed from cells, replaced with assay medium and microplate was incubated 45 minutes at 37°C without CO2. Extracellular flux cartridge was prepared adding 0.5μM rotenone/antimycin A in port A and 50mM 2-Deoxyglucose in port B diluted in Xfp assay medium and the cartridge was calibrated by Seahorse XFp analyser setting Glycolysis Test program. When the calibration was finished, cell microplate was inserted in a Seahorse XFp analyser starting the analysis.

### RNA extraction and qRT-PCR

3.0×10^5^ cells per well were seeded in a 6-well plate and incubated overnight. The day after, cells were starved and lysed with β-mercaptoethanol and RNA was extracted using Illustra RNAspin Mini RNA Isolation kit (GE-Healthcare, Chicago, IL, USA) following the kit protocol. Then, RNA was retro-transcribed using the RevertAid RT Kit (Thermo Fisher Scientific). Finally, RT-qPCR was performed using 25ng and 2x qPCR SyGreen (PCR Biosystem, London, England) in Cfx96™ Real-Time System or Cfx384™ Real-Time System Thermocyclers (Bio-Rad). The sequence of all primers used is reported in **Table S2**.

### Digital droplet PCR

Digital droplet PCR was performed based on the protocol developed by Bio-Rad. Briefly, 1×10^5^ cells were seeded in 6-well plate and grown for 1 day. Next cells were detached, counted and solubilized in DireCtQuant 100ST solubilization reagent (DireCtQuant, Lleida, Spain) at the concentration of 1000 cells/μL, incubating for 3 minutes at 90°C with shaking at 750 rpm and then centrifuged for 1 minute at 10000 g. Lysates were obtained by incubation for 3 minutes at 90°C under shaking at 750 rpm and then centrifuged for 1 minute at 10000 g. Lysate was diluted 1:4 in solubilization reagent and for every target to analyze, PCR mix was prepared comprising 2μL of diluted sample, forward and reverse primers at 10μM final concentration in a final volume of 95μL. PCR mix without lysate was used as negative control. Then, 10.5μL of each PCR reaction was mixed with 11μL of 2xQX200 ddPCR EvaGreen SuperMix (Bio-Rad) and 0.5μL of HindIII restriction enzyme (NEB, Ipswich, MA, USA). Sample DNA was digested at 37°C for 15 minutes and loaded in ddPCR DG8™ Cartridge with QX200™ droplet generation oil (Bio-Rad). Droplets were made using QX200™ droplet generator (Bio-Rad) and then loaded in a 96-well PCR plate. Sample DNA was amplified and finally, droplets with amplified DNA were analyzed using the QX200™ droplet reader and QuantaSoft 1.7.4 (Bio-Rad).

### RNA Immunoprecipitation

HCT116 cells were cultured in standard medium in P150 plates till they reached ~80% confluence. ~10^7^ cells were lysed in 1mL of Lysis buffer (100mM KCl, 5mM MgCl2, 10mM HEPES pH 7, 0.5% NP-40, 1mM DTT, 1U/ul RNase Inhibitors, 1X Protease Inhibitor Cocktail) using a scraper. Lysates were transferred in a falcon tube, placed for at least two hours at −80°C, and centrifuged at 10000 rpm for 30 minutes. Supernatants were collected in a new tube. Dynabeads ProteinA or ProteinG (depending on the antibody species, Thermo Fisher scientific) were prepared by washing them twice with NT2 Buffer (50mM Tris-HCl pH7.4, 150mM NaCl, 1mM MgCl2; 0.05% NP40) and resuspended in NT2 buffer. Beads were distributed in different tubes, supplemented with twice their initial volume of NT2 Buffer. The DHX30 specific antibody (5 μg -A302-218A, Bethyl) or IgGs were added to the beads and incubated for 2 hours on a rotating wheel at 4°C. Lysates were pre-cleared by adding a mix containing ProteinA and ProteinG Dynabeads in equal amounts and incubating for 1 hour at 4°C on a wheel. After placing the tubes on a magnet, supernatants were collected and 1% of their volume was used as input to be directly extracted with TRIzol. The remaining supernatant was added to the antibody-coated beads and incubated overnight on a wheel at 4°C. The day after, beads were washed with 1ml NT2 buffer for 10 minutes on a wheel at 4°C. Three additional washes were performed again with 1ml NT2 Buffer supplemented with 0.1% Urea + 50mM NaCl (10 minutes each at 4°C on a wheel). Beads were washed one more time in 500μl of NT2 buffer, 50μl were collected for WB analysis and the remaining supernatant was discarded. RNA was extracted by adding TRIzol to the beads, according to the manufacturer’s protocol. The RNA pellets were resuspended in 15μl DEPC water and cDNAs were synthesized using the RevertAid First Strand cDNA Synthesis Kit (Thermo Fisher Scientific).

### Cytoplasm-Mitochondria fractionation

For cytoplasm-mitochondria fractionation, 2×10^6^ cells were seeded in a P150 Petri dish and incubated for 2 day. Then, they were detached using trypsin and re-suspended in 750μL of Mitochondrial Isolation Buffer (MIB) (0.32M Sucrose, 1mM EGTA pH 8.0, 20mM Tris-HCl pH 7.2) with 0.1% fatty acid-free BSA. Cells were homogenized in a Potter-Elvehjem homogenizer (DWK Life Sciences, Mainz, Germany) applying more than 60 strokes. Homogenate (A) was centrifuged for 5 minutes at 1000g at 4°C and supernatant was collected. Cell debris were resuspended in 750μL of MIB + BSA and re-homogenized with more than 20 strokes. New homogenate (B) was centrifuged for 5 minutes at 1000g at 4°C and the supernatant was collected and pooled with supernatant A. Two mixed supernatants (A+B) were centrifuged at 12000g for 10 minutes at 4°C to pellet the mitochondria. Supernatant was discarded and mitochondria were washed twice with 1mL of MIB + BSA and once with 1mL of MIB, centrifuging the mitochondrial pellet every time at 12000g for 10 minutes at 4°C. Finally, supernatant was discarded and mitochondria were lysed in RIPA buffer and mitochondrial proteins were quantified by BCA assay.

### Apoptosis assay

Apoptosis analysis was based on FITC Annexin-V Apoptosis Detection Kit protocol (BD Biosciences, San Jose, CA, USA). Briefly, 3×10^5^ cells per well were seeded in a 6-well plate, incubated for one day and then treated with drugs under investigation for 48 hours. 0.1% DMSO was used as negative control. Cells were detached using trypsin and washed twice with PBS 1X. Then, 1.5×10^5^ cells were resuspended in 100μL of Binding Buffer 1X, stained with FITC-Annexin V and PI and incubated for 15 minutes at room temperature in the dark. Finally, samples were analyzed by FACSCanto™ flow cytometer and BD Diva Software 6.1.3. Unstained, PI-only and FITC-only stained samples were used to set the cytometer’s parameters.

### Cell cycle analysis

HCT shNT or shDHX30 were seeded in 6-well plates at a concentration of 3×10^5^ cell/well. 48 hours after seeding, cells were trypsinized, washed with 1X PBS and counted for subsequent staining according to BD Cycletest™ Plus DNA Kit protocol. Briefly, 5×10^5^cells were centrifuged, superantant was removed and 250μl of Solution A were added. Cells were resuspended and incubated for 10 minutes before adding 200 μl of Solution B. After 10 minutes of incubation, 200 μl of ice-cold Solution C were added to the samples and cells were incubated for 10 minutes at 4°C in the dark before analysis with FACS CantoA (BD). The percentage of cells in each stage of the cell cycle (G1, S or G2) was computed using the ModFIT LT 4.0 software (Verity Software House).

### Exploration of TGCA data using the GEPIA web resource

The GEPIA API [17,18] was used to extract correlations between DHX30 and mitoribosomal genes (either as individual genes or for the 14-genes signature) in all available TCGA tumor datasets (correlation mode). The same analysis was performed to correlate cytoplasmic ribosome and mitochondrial ribosome genes expression, on TCGA tumor samples, matched TCGA normal samples, and GTEX healthy tissues RNA-seq data. Survival analysis was performed with the same tool and in the same datasets, using the survival mode. Only results within the p-value threshold of 0.05 were considered, with the others being set to R=0 and HR=1 in the heatmap for visualization purposes.

### Statistical analysis

Data are presented as mean ± SD of three independent biological experiments, unless stated otherwise. GraphPad Prism 5 or 9 software (GraphPad software, La Jolla, CA, USA) were used and either a two-way anova with Tukey’s multiple comparison test or a two-tailed Student’s t-test was performed, unless specified otherwise.

## Data availability

The RNA-seq datasets are available in GEO with ID GSE95024 and GSE154065.

## Acknowledgements

We thank Pamela Gatto, Viktoryia Sirarovich and Michael Pancher of the HTS CIBIO core facility for assistance with droplet digital PCR experiments, high-content as well as impedance-based proliferation assays. We thank Nicoletta di Bernardo for technical assistance for the apoptosis assays. This work was supported by the Italian Association for Cancer Research (Fondazione AIRC, IG grant #18985 to AI). JJD laboratory is supported by grants by the Institut National du Cancer (INCa, grant PRT-K program n°2018-024 EMT-CoNCEPT) and Agence Nationale de la Recherche (ANR, grant RiboCard ANR-18-CE11-0020, and grant ActiMeth ANR-19-CE12-0004).

## Authors contributions

B.B. perfomed most experiments and wrote the first draft of the paper. A.R., D.R., S.G., A.P., F.B., and, A.G. helped in some experimental procedures. F.C. and J-J.D. helped to design some experiments. E.D. performed all data analysis. E.D. and A.I. designed the study. A.I. co-wrote the first draft of the manuscript.

---

**Table S1: RNA-seq Fold Change data and results of GSEA analysis for the comparison between HCT_shDHX30 and HCT_shNT.**

A), B) ID, gene symbol, fold change, F value, P value and FDR based on RNA-seq data of (A) polysomal RNA, (B) total RNA. GSEA data for polysomal (C) or total (D) RNA-seq data.

**Table S2: siRNAs, FISH probe, and primers.**

Contains information on all primers used in qRT-PCR experiments.

**Table S3: List of primary and secondary antibodies**

