## Supplemental Information for "DHX30 coordinates cytoplasmic translation and mitochondrial function contributing to cancer cell survival"

**Bartolomeo Bosco *et al.***

**SUPPLEMENTARY FIGURES & LEGENDS**

A

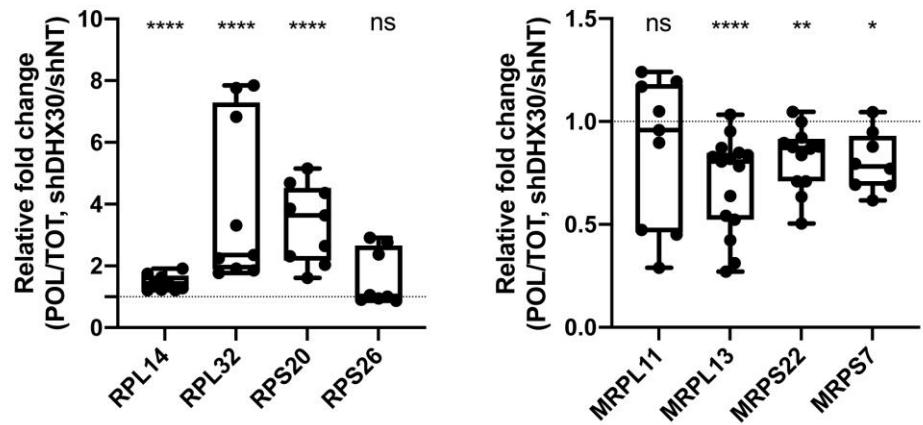

B

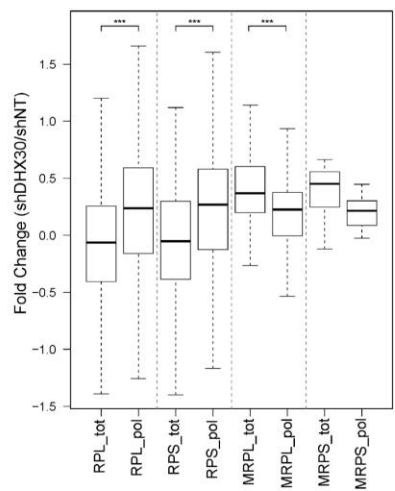

C

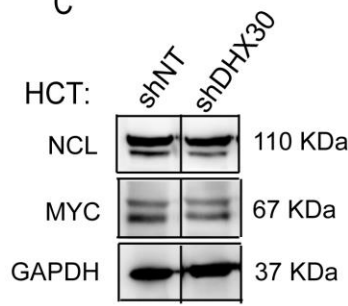

D

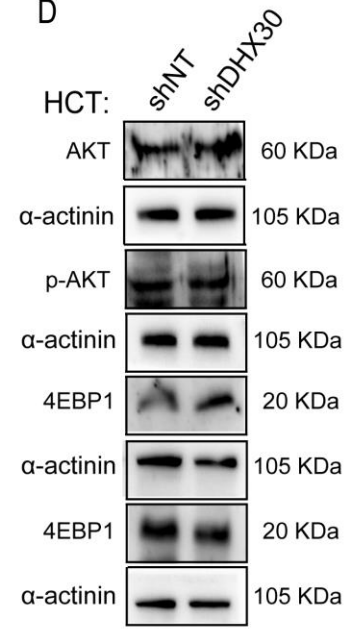

E

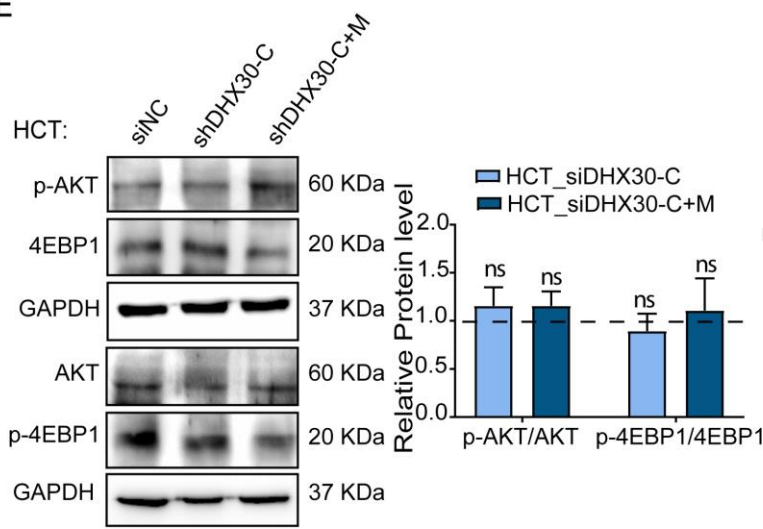

F

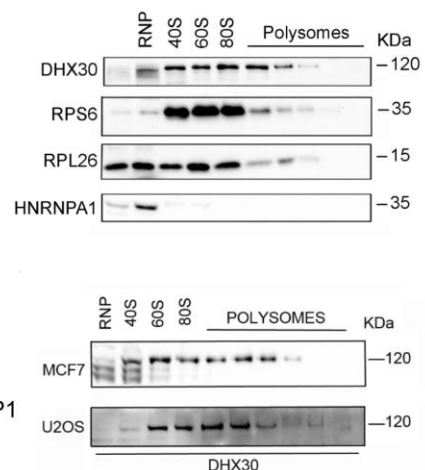

**Figure S1. Relative expression of ribosomal and mitoribosomal transcripts, status of MYC and mTOR pathway and association of DHX30 with ribosomes. Related to Figure 1.**

A) For both panels, relative translation efficiency obtained from RT-qPCR data for the indicated transcripts starting from total or polysomal RNA of HCT116 control shDHX30 cells. Data are plotted as the relative fold change in the comparison between polysomal and total RNA first, for shDHX30 over shNT. n=2 biological replicates each with three technical replicates. YWHAZ and B2M were used as references. **\*\*p** < 0.01, two-tailed Mann Whitney test. **B)** DHX30 depletion leads to slight changes in the expression of ribosomal protein genes. Box plot of the expression fold changes in both total and polysomal RNAs for the indicated transcript groups in HCT\_shDHX30 relative to the HCT116\_shNT control. **\*\*\*p** < 0.001. **C)** Stable DHX30 silencing does not lead to alteration in the expression of c-MYC and of its target nucleolin (NCL). A representative western blot image is shown. **D** and **E)** Neither stable nor transient silencing of DHX30 leads to activation of the MTOR pathway, based on the relative phosphorylation of AKT and 4EBP1. See text and Figure S2 for details on the transient DHX30 silencing. Left panels in **E)**, representative blot images. Right panels, relative quantification of the ratio between phospho and total AKT or 4EBP1 and in the comparison between DHX30 silenced cells and control (siNC) cells, respectively. **F)** Top panels, western blot to visualize DHX30 distribution in the different fractions of HC116 cells after polysome profiling obtained by a linear 15-50% sucrose gradient. RPS6, RPL26, and HNRNPA1 were used as controls. Bottom panels, DHX30 protein is associated with ribosomal subunits and polysomes in MCF-7 and U2OS cells. Proteins were extracted from each fraction and subjected to western blot analysis using an anti-DHX30 antibody for immunodetection.

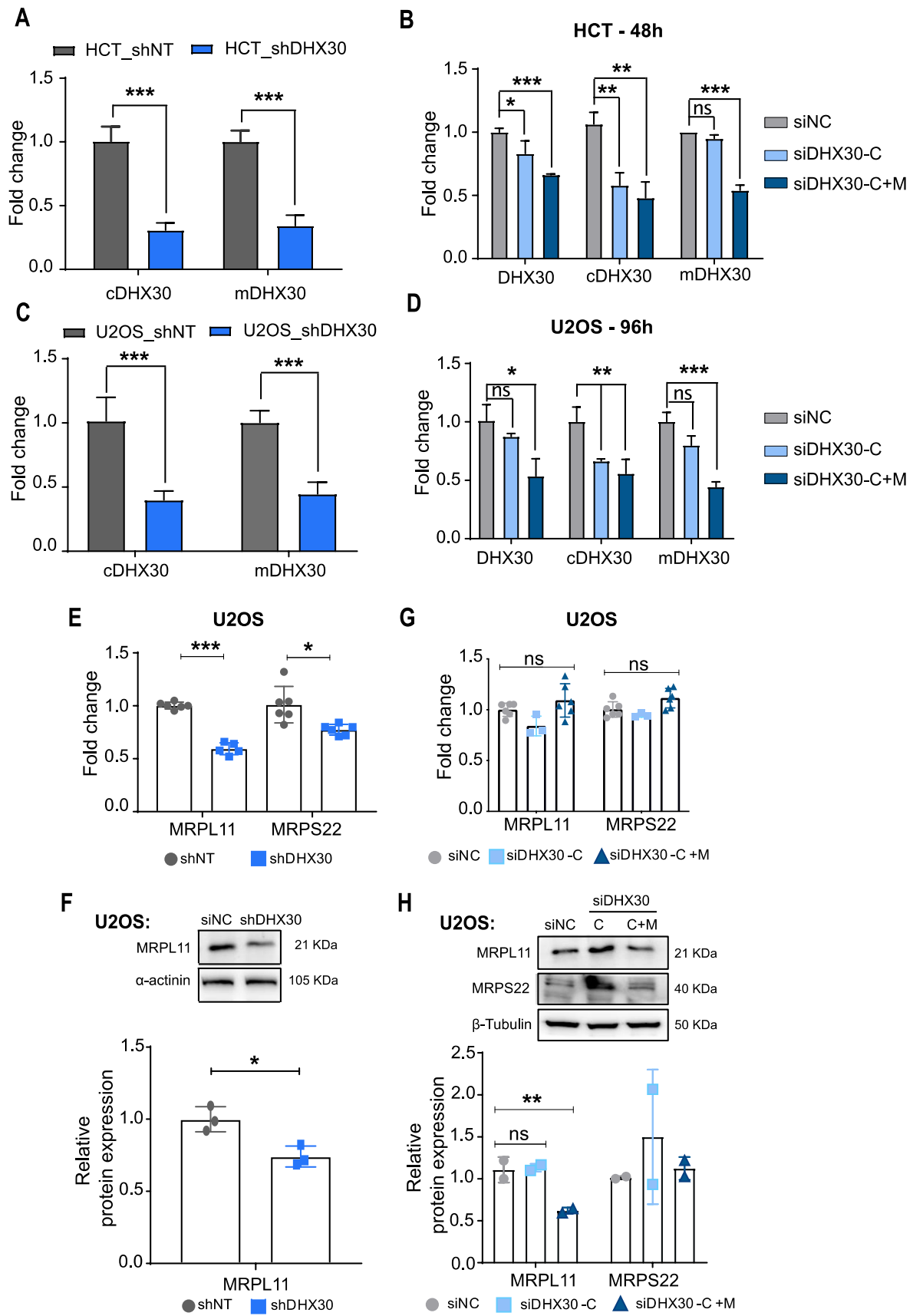

**Figure S2. Efficacy of both stable and transient DHX30 silencing in HCT116 and U2OS cells and impact of DHX30 depletion on nuclear encoded mitoribosome components. Related to Figure 2 & 3.**

**A)** Relative mRNA levels of cytoplasmic (cDHX30) and mitochondrial (mDHX30) DHX30 transcript variants in HCT116\_shDHX30 compared to the shNT control clone. Data are mean ± SD (n=3); \*\*\*p < 0.001. **B)** qRT-PCR to verify the transient silencing of DHX30 in HCT116 using the indicated siRNAs for 48 hours. PCR primers annealing to: (i) a portion of the coding sequence that is present in all transcript variants (DHX30); (ii) the first exon specific of cytoplasmic DHX30 (cDHX30) or (iii) the alternative first exon specific of mitochondrial DHX30 (mDHX30) were used. Data are mean ± SD (n=3); \*p < 0.05, \*\*p < 0.01; \*\*\*p < 0.001. **C)** Same as A, but for U2OS\_shNT and U2OS\_shDHX30 clones. Data are mean ± SD (n=3); \*\*\*p < 0.001. **D)** Same as B, except that U2OS cells were transiently transfected and qPCR was performed using RNA extracted 96 hours after the transfection. **E)** qRT-PCR of MRPL11 and MRPS22 in U2OS\_shDHX30 cells relative to the U2OS\_shNT control (set to 1). Average, standard deviation, and individual data points are shown (\*p < 0.05; \*\*\*p < 0.001). **F)** (Upper panel) representative western blot of MRPL11 in U2OS\_shNT or U2OS\_shDHX30 cells. (Lower panel) Relative protein quantification; mean, SD, and individual points are shown; \*p < 0.05. **G)** qRT-PCR of MRPL11 and MRPS22 in U2OS cells transiently silenced for cytoplasmic DHX30 (siDHX30-C) or for both cytoplasmic and mitochondrial variants (siDHX30-C+M) for 96 hours. Data are compared to the siRNA negative control (siNC) and are mean ± SD (n=3); individual data points are also shown. **H)** (Upper panel) representative western blot of MRPL11 and MRPS22 in U2OS silenced transiently as in A). (Lower panel) Relative protein quantifications; mean, confidence interval, and individual points are shown; \*\*p < 0.01.

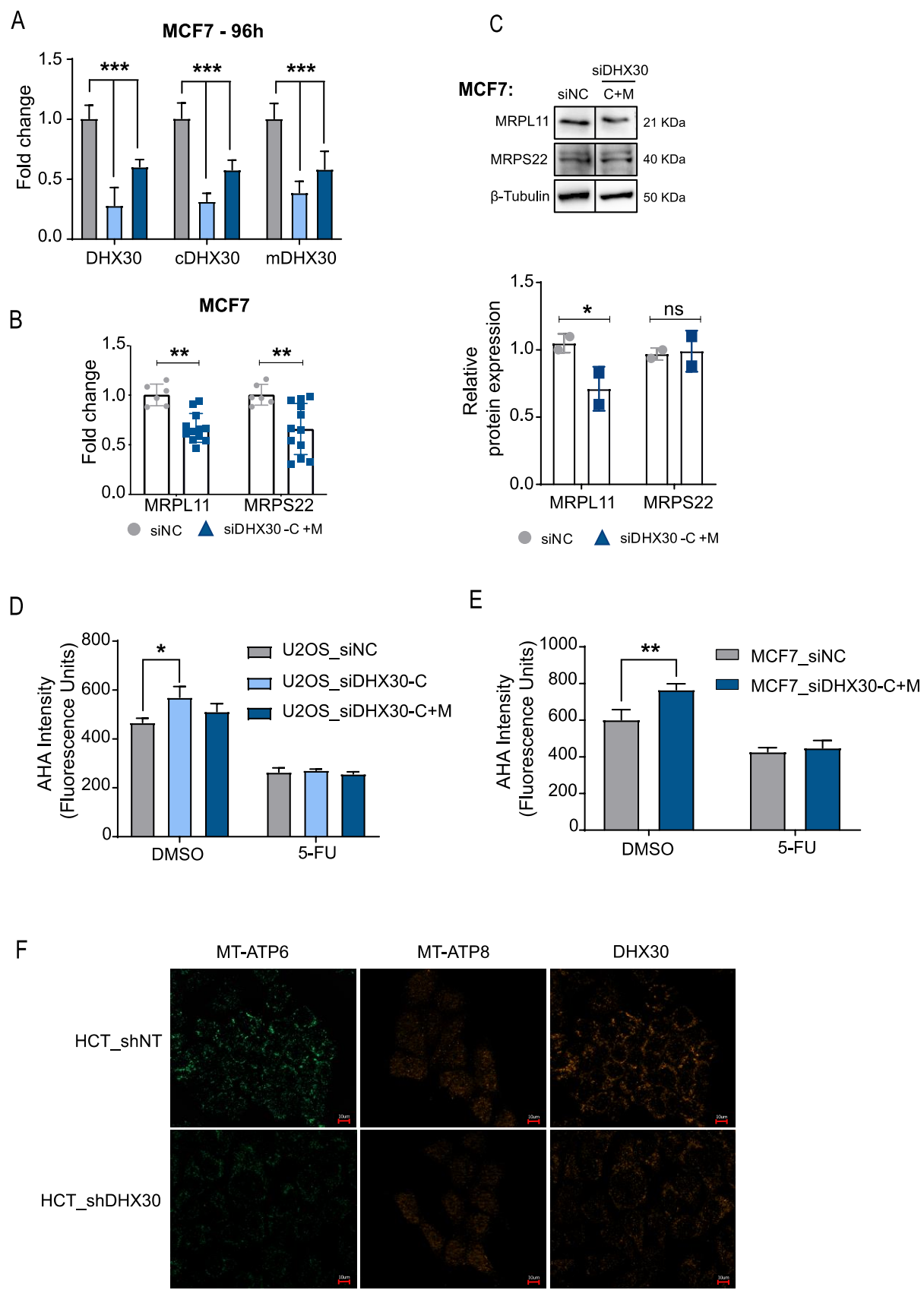

**Figure S3. Impact of DHX30 depletion on nuclear encoded mitoribosome components in MCF7 and U2OS, and on mitochondrial OXPHOS components in HCT116 cells. Related to Figure 2.**

**A)** qRT-PCR to verify the efficacy of the transient silencing of DHX30 MCF-7. Samples were collected for processing 96 hours after silencing with corresponding siRNA. Data are mean ± SD (n=3); \*\*\*p < 0.001. Surprisingly siDHX30-C was not isoform-specific in MCF7 and, hence, removed from subsequent tests. The apparent lack of specificity remains to be investigated. **B)** qRT-PCR of MRPL11 and MRPS22 transcripts in MCF7 cells transiently silenced using siDHX30-C+M for 96 hours. Data are compared to the siRNA negative control (siNC) and are mean ± SD (n=3); individual data points are also shown; \*\*p < 0.01. **C)** (Upper panel) representative western blot of MRPL11 and MRPS22 in MCF7 silenced transiently for both cytoplasmic and mitochondrial variants (MCF7\_siDHX30-C+M). (Lower panel) Relative protein quantifications; mean, confidence interval, and individual points are shown; \*p < 0.05. **D)** Analysis of global translation based on L-azidohomoalanine (AHA) fluorescence intensity measured from cytoplasmic proteins in U2OS cells transiently transfected as indicated for 96 hours. 5-Fluorouracil treatment was used as a control leading to reduced translation. Data are mean ± SD (n=3); \*p < 0.05. **E)** Same as D, but for transiently transfected MCF7 cells; \*\*p < 0.01. **F)** Representative images of one of three independent immunofluorescence experiments to visualize DHX30, MT-ATP8 (red), or MT-ATP6 (green) expression in HCT116\_shDHX30 and HCT116\_shNT.

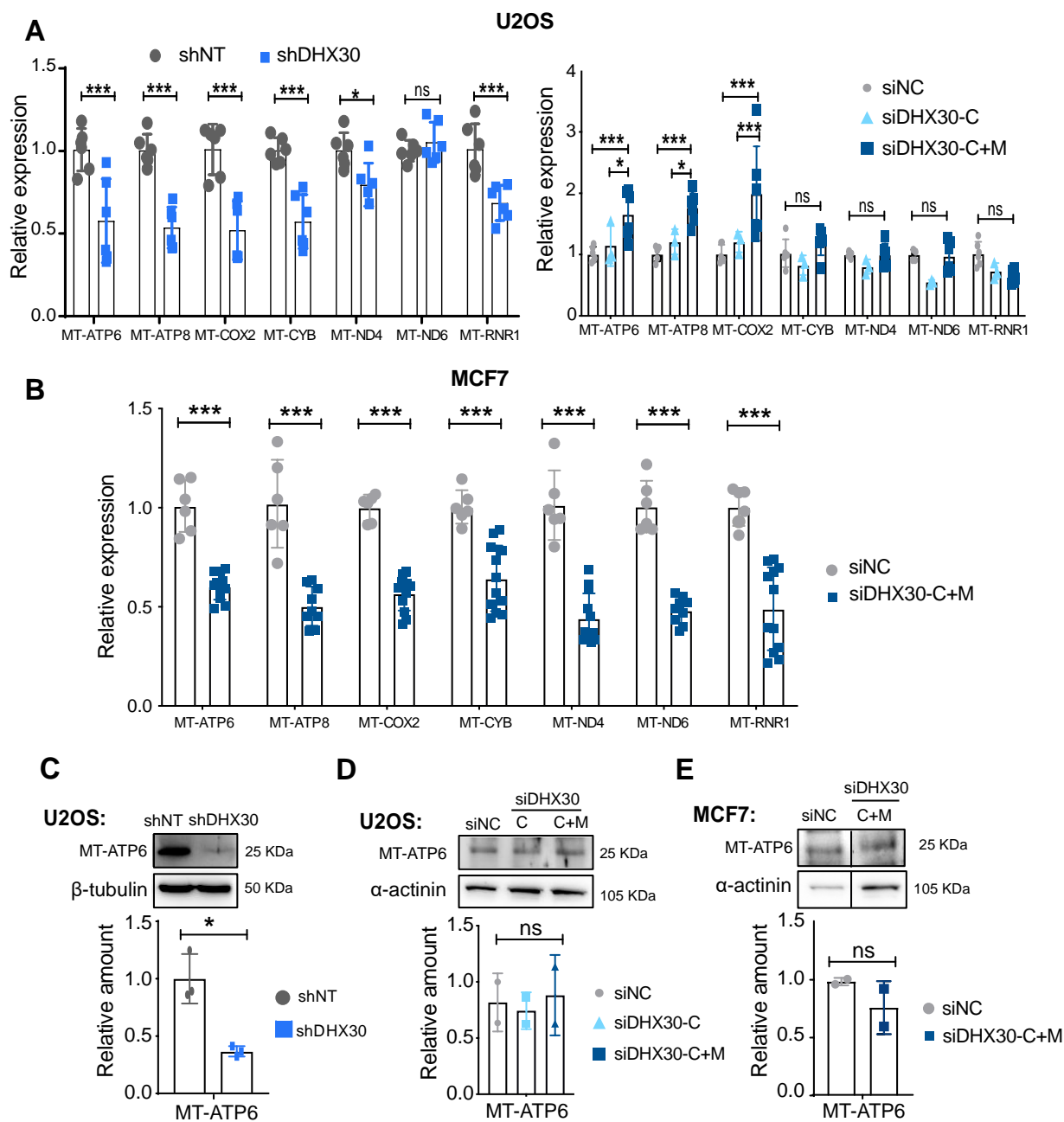

**Figure S4. Impact of DHX30 depletion on mitochondrially encoded OXPHOS components and on proliferation in MCF7 and U2OS cells. Related to Figure 3 and Figure 4.**

**A)** (left panel) qRT-PCR of selected mitochondria-encoded genes in U2OS<sub>shDHX30</sub> relative to the U2OS<sub>shNT</sub> control (set to 1); (right panel) U2OS transiently silenced for cytoplasmic DHX30 (siDHX30-C) or for both cytoplasmic and mitochondrial variants (siDHX30-C+M) for 96 hours. Average, standard deviations, and individual data points are shown (\*$p < 0.05$; \*\*\*$p < 0.001$). For transient silencing, data are compared to the siRNA negative control (siNC). **B)** qRT-PCR of selected mitochondria-encoded genes in MCF7 cells transiently silenced for both cytoplasmic and mitochondrial DHX30 (siDHX30-C+M) for 96 hours. Data are compared to the siRNA negative control (siNC). Mean and individual points are shown ($n=2$ biological replicates); \*\*\*$p < 0.001$. **C-E)** (Upper panels) representative western blot of MT-ATP6 in U2OS<sub>shNT</sub> or <sub>shDHX30</sub>, and in U2OS or MCF7 transiently silenced as in A) or B). (Lower panels) Relative protein quantification of MT-ATP6. Mean, SD, and individual points are shown ($n=3$ for <sub>shDHX30</sub> clone; $n=2$ for siRNA experiments); \*$p < 0.05$.

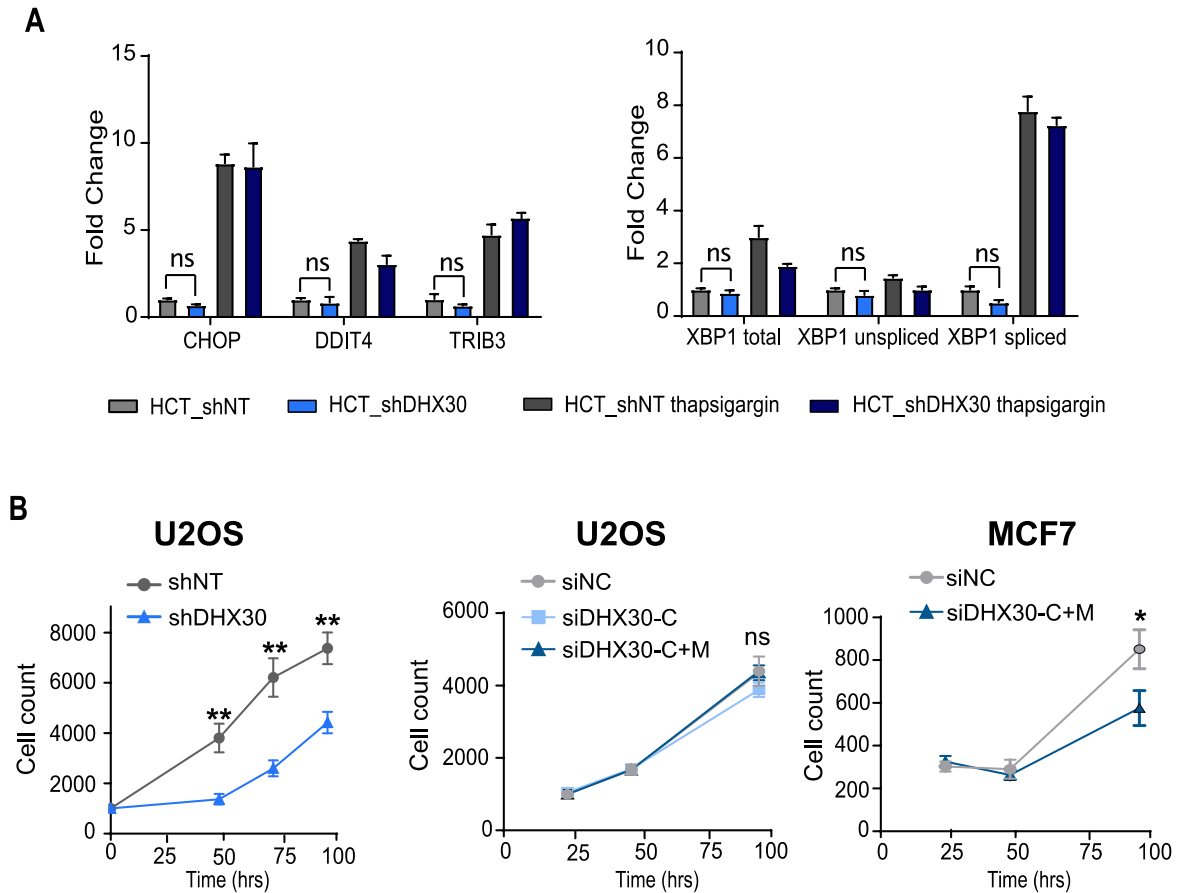

**Figure S5. Impact of DHX30 depletion on the ER-stress response genes and on cell proliferation. Related to Figure 5.**

**A)** RT-qPCR data for the indicated ER-stress response genes. Bars plot the average fold change and the SD of three replicates. No significant differences were observed in the comparison between HCT116<sub>shNT</sub> and <sub>shDHX30</sub> cells. Treatment with Thapsigargin (100 nM for 24h) was included as a positive control for ER-stress. Also in this case, while gene expression of XBP1 splicing increased, there was no apparent effect associated with DHX30 depletion. **B)** Left panel, relative cell proliferation measured by high-content microscopy in digital phase contrast in U2OS<sub>shNT</sub> or U2OS<sub>shDHX30</sub>. U2OS (center panel), or MCF7 (right panel) transiently silenced for DHX30 variants. Data are mean ± SD (n=2 biological replicates each with three independent wells); \*p < 0.05.

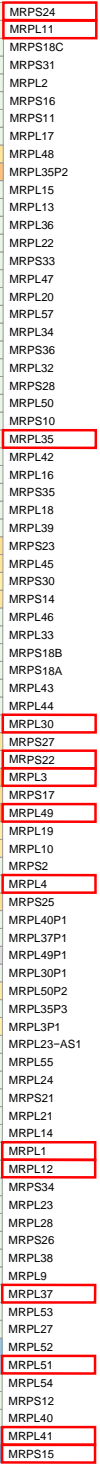

Pearson correlation between the expression of DHX30 and each mitoribosomal protein transcript in cancer samples from TCGA. Data were extracted from the GEPIA web server (see methods for details).

Unsupervised clustering revealed two major clusters based on the level of gene expression correlation and two main clusters among cancer types. Boxed in red are the names of the fourteen transcripts that are candidate DHX30 direct targets (see text for details and Figure 6).

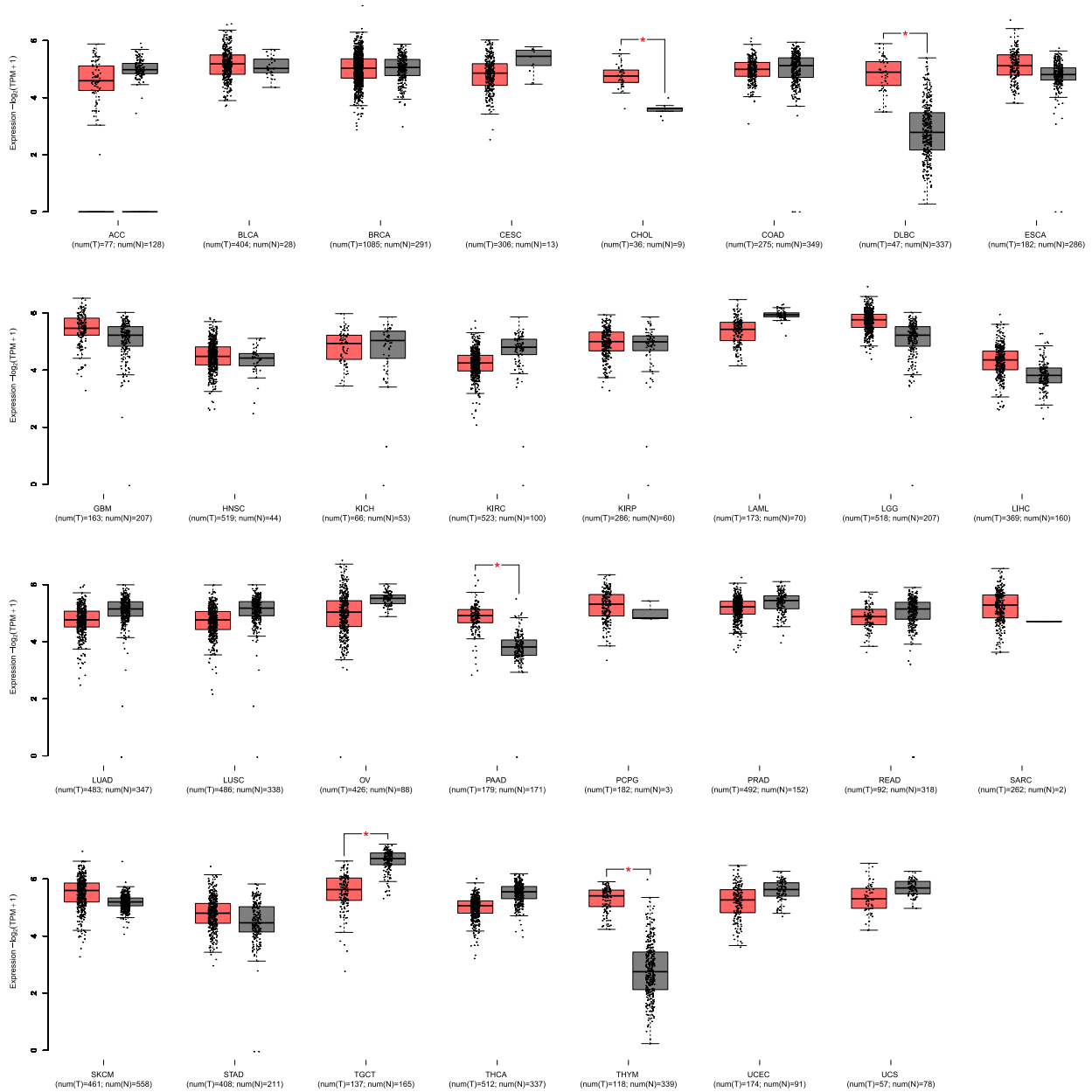

**Figure S7. Expression of DHX30 in TCGA tumor types and matched healthy tissues. Related to Figure 6.**

Each boxplot pair displays the expression levels of DHX30 in TCGA tumor samples (red box) and in the corresponding healthy tissue samples (grey box, composed of TCGA normal samples and GTEX samples of the same tissue). Expression is displayed in $\log_2(\text{tags per million sequenced reads} + 1)$. Plots were obtained with GEPIA2. Significance of the difference between tumor and normal samples is indicated by a red asterisk and represents a $q\text{-value} \leq 0.01$ and a $|\log_2\text{FC}| > 1$ as computed in GEPIA2 by the ANOVA method.

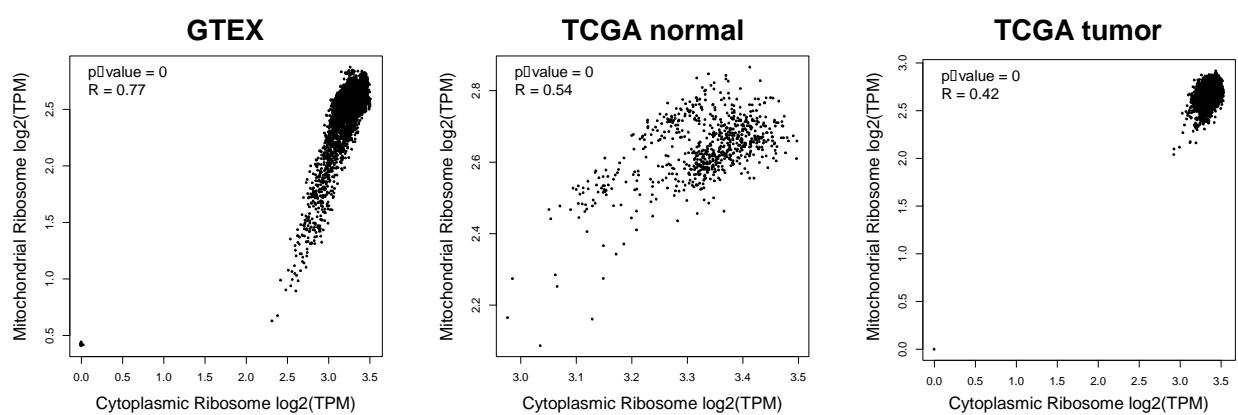

**Figure S8. Expression correlation of cytoplasmic and mitochondrial ribosome component genes in TCGA and GTEX samples. Related to Figure 6.**

The three scatterplots display the Pearson correlation ($R$ and $p$-value indicated in the upper left corner of each plot) between the expression (indicated as $\log_2(\text{tags per million sequenced reads})$) of cytoplasmic ribosome component genes (X axis) and mitochondrial ribosome component genes (Y axis). The leftmost plot shows the correlation in GTEX healthy tissues samples, the center plot presents the correlation in TCGA normal samples, and the rightmost plot shows the correlation in TCGA tumor samples. Plots were obtained with GEPIA2.

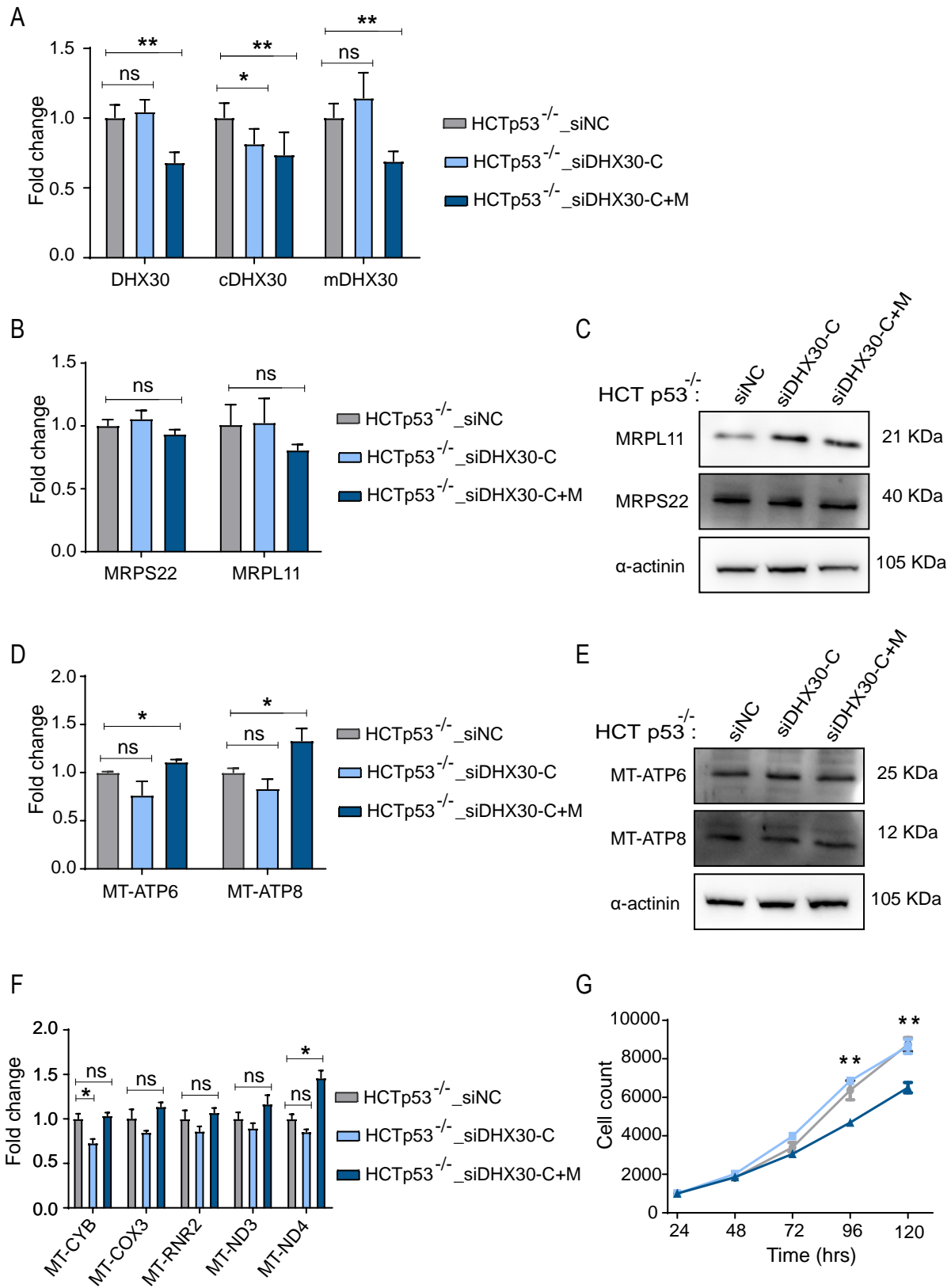

**Figure S9. Silencing of DHX30 does not affect mitoribosome and OXPHOS components in HCT116 when p53 is knock-out. Related to Figures 2-5.**

**A)** qRT-PCR to verify the transient silencing of DHX30 in HCT116 p53<sup>-/-</sup> using primers annealing to: (i) a portion of the coding sequence that is present in all transcript variants (tDHX30); (ii) the first exon specific of cytoplasmic DHX30 (cDHX30) or (iii) the alternative first exon specific of mitochondrial DHX30

(mDHX30). Experiment was performed 96 hours post silencing. **B), C)** Relative mRNA **B)** and protein **C)** levels of MRPL11 and MRPS22 in HCT116 p53<sup>-/-</sup> transiently silenced as shown in **A)** and compared to the siNC control. α-actinin was used as loading control. Immunoblots represent one of three independent experiments. **D), E)** Relative mRNA **D)** and protein **E)** levels of MT-ATP6 and MT-ATP8 in HCT116 p53<sup>-/-</sup> transiently silenced as shown in **A)** and compared to the siNC control. α-actinin was used as loading control. Immunoblots represent one of three independent experiments. **F)** RT-qPCR to study some of mitochondrially encoded OXPHOS components in HCT116 p53<sup>-/-</sup> transiently silenced for DHX30 variants compared to the siNC control. **G)** Relative cell proliferation of HCT116 p53<sup>-/-</sup> cells measured by high-content microscopy in digital phase contrast and compared to siNC control at indicated time points. For all panels, data are mean ± SD (n=3); \*p < 0.05; \*\*p < 0.01.

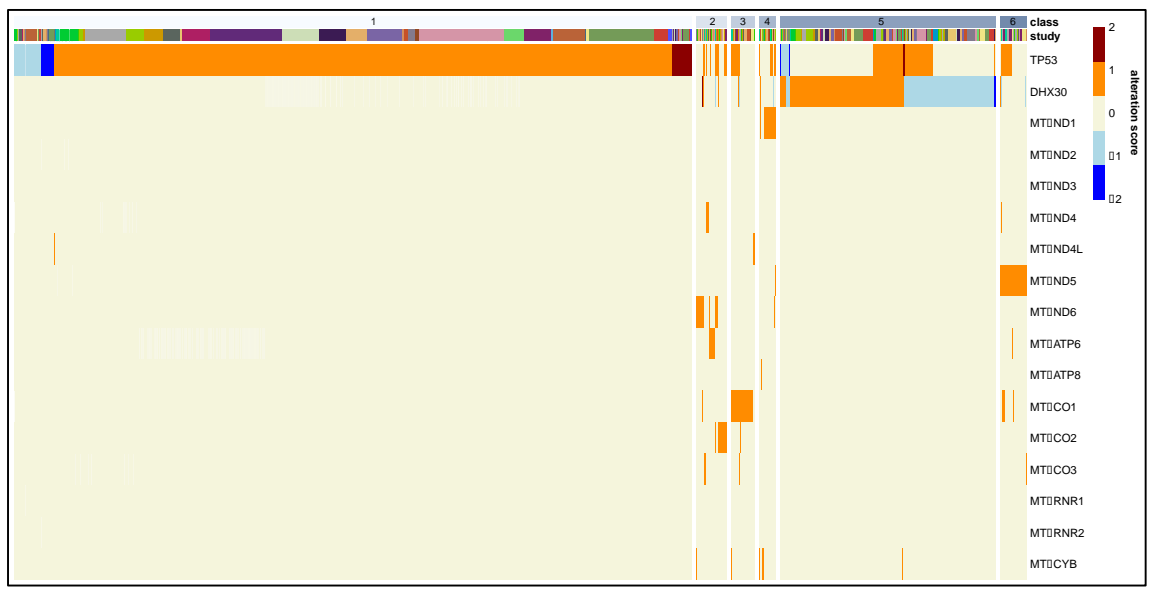

**Figure S10. DHX30 copy number, mutation, and expression alterations in human cancer. Related to Figure 6.**

Pan cancer data from TCGA was interrogated for copy number alterations, mutations and expression alterations of DHX30, TP53 and mitochondria-encoded genes. A low frequency of DHX30 alteration was apparent and did not appear to be associated with a specific cancer type. No apparent correlation between DHX30 and TP53 alteration status was apparent. Instead, although limited by the low number of occurrences, alterations in mitochondrial genome and DHX30 CNVs seemed to be mutually exclusive.
